# Cerebellar Purkinje cells change dendritic architecture during primate evolution

**DOI:** 10.64898/2026.06.07.728033

**Authors:** Aurora Ferrell, Silas E. Busch, Madison Hillegas, Chet C. Sherwood, Christian Hansel

## Abstract

Vertebrate evolution has driven adaptive remodeling of brain regions impacted by changing sensorimotor demands. The cerebellum—a hindbrain area mediating associative learning and predictive coding—is thought to participate in diverse motor and non-motor behaviors across vertebrate species by duplicating and repurposing a conserved cortical circuit. Yet, few studies systematically compare circuit architecture across cerebellar evolution. We recently found that human Purkinje cells (PCs, the principal neuron of the cerebellar cortex) almost universally have multiple primary dendrites, a structural motif that confers distinct signaling properties, as shown by experiments in mice where this motif is present but less common. Seeking evolutionary insight, we developed a framework to parameterize and compare PC morphology across 11 simiiform (anthropoid) primates representing 40 million years of evolution, and mice as an outgroup. Dendritic architecture shifts profoundly from single primary dendrites with vertical orientation in mice and monkeys to multiple horizontally oriented primary dendrites in apes and, particularly pronounced, in humans. Increasing dendritic compartmentalization in the human lineage is produced by monotonic, stepwise, or human-specific trends across individual morphological traits. PC morphology is high-dimensional, with most features varying widely and independently, yet clade identity can be readily predicted from multivariate morphospace profiles. Phylogenetic patterns are not well explained by tissue foliation or allometric scaling, supporting the hypothesis that morphological variation is constrained by functional demands. Echoing a core principle of brain evolution that selection pressure drives fine-tuning of circuit elements rather than total circuit rearrangement, PC dendrite morphology may serve as a key node for cerebellar adaptation amid a conserved circuit architecture.

## Introduction

The cerebellum plays a conserved role in detecting and correcting errors in sensorimotor, affective, and cognitive domains across vertebrate species^1–4^. These functions are achieved by comparing ongoing contextual information with learned and stored internal models^5–7^. This comparison involves the integration of two distinct excitatory pathways by a single neuron type, the Purkinje cell (PC), which then produces the sole output of the cerebellar cortex^8^. Information about ongoing internal and external state arrives on the PC dendritic arbor in the form of thousands of parallel fiber (PF) axons^9,10^, making PCs the most input-dense neurons in the brain^11^. Cerebellar learning and the processing of unexpected stimuli occurs through bidirectional plasticity of PF synapses^12–14^. The direction of PF plasticity is mediated by the relative timing of PFs and ‘teaching’ signals provided by climbing fibers (CFs), a coincidence detection algorithm with a functional signature shared by neural circuits across brain areas, species, and neural network models^15–19^. In the classic understanding of cerebellar histology, each PC receives input from only one CF—the winner of competitive, activity-dependent CF pruning during early postnatal development^20^—rendering each PC into a single functional compartment^12,21,22^.

Since its earliest anatomical^23^ and physiological^5,9,21,24^ characterization, this circuit has been regarded as highly uniform^25,26^. However, recent work shows that PCs, more so than their neighboring cell types in the cerebellum, are heterogeneous at the molecular, physiological, and morphological levels. Patterned zebrin and parvalbumin expression in rodents produces PC subtypes with distinct firing patterns and plasticity rules^27–30^. There is pronounced heterogeneity in developmental subtype generation between PCs as described in mice and humans, but not other cerebellar cell-types^31–33^. Comparative immunolabeling is suggestive of variation in PC morphology across mammalian species^34,35^, which is recently becoming systematically characterized across cerebellar regions of humans and mice^11,36–38^. These findings suggest that the otherwise uniform cerebellar circuit may exhibit heterogeneity at the level of PCs.

In adult mice, roughly a quarter of PCs with multiple primary dendrites—emerging either from the soma or a proximal bifurcation of a single trunk—receive multiple CF inputs with distinct receptive fields^36^. This multidendritic motif is uncommon in mice and nearly universal in humans, yet, both species exhibit conserved regional variation whereby multi-dendritic PCs are more common in the posterior hemisphere, which integrates multi-modal information in mice^36,39–45^ and humans^46–49^. Computational models of human PCs can host 6-7x more input combinations than mouse PCs owing to differences in dendritic compartmentalization and size^37^. PC primary dendrite morphology may thus serve as a determinant of region-specific, and possibly species-specific function. We currently lack systematic comparative anatomical data for PCs^35^ and the associated circuit architecture, leaving open the question: is PC morphology a critical node for evolutionary adaptation?

We developed a comparative, parametric analysis of PC morphology across 11 simiiform (anthropoid) primate species—representing major primate clades, platyrrhine and cercopithecid monkeys, hylobatid and hominid apes, and excluding only lemuriforms (lemurs and lorises) and tarsiiforms (tarsiers) clades—spanning ∼40 million years of primate evolution. We also examined PC morphology in mice as an outgroup comparison, representing a rodent species that diverged from primates ∼85 million years ago. Though simiiforms represent a small fraction of vertebrate and mammalian evolution (∼530 and 250 million years, respectively), we observed substantial variation in their PC morphology, which become increasingly large and compartmentalized along the human evolutionary trajectory. Many morphological features are highly variable and do not covary, consistent with a high-dimensional system and possibly reflecting independent and permissive developmental regulation. Despite this variability, clade-specific patterns of adaptation emerge from PC multivariate morphospace profiles. Morphological trends are poorly explained by tissue foliation and allometric scaling. Collectively, we show that PC morphology represents a key node for adaptation in the geometrically precise cerebellar cortical circuit. This supports the conclusion that the cerebellum is not architecturally uniform and may vary to support distinct functional needs, positioning it as a powerful inroad to understanding the evolution of vertebrate neural circuits.

## Results

### PC demographics shift across primate evolution

We recently found that human PCs are morphologically distinct from those of mice^11,36^, but whether those differences arose uniquely in humans or evolved gradually as the species diverged is not known. To address this question directly, we asked how human PC morphology compares with that of our closest relatives: simiiform primates. We obtained cerebellar tissue from 3 mice, as a reference to earlier divergence in mammalian evolution, and from 26 individuals across 11 species (Table S1, Figure 1A), including platyrrhine monkeys (common and white-faced marmoset and black-handed spider monkey), cercopithecid monkeys (rhesus macaque and olive baboon), hylobatids (northern white-cheeked gibbon and siamang), and five species of hominid apes (Sumatran and Bornean orangutan, western lowland gorilla, chimpanzee, bonobo, and human) and immunolabeled PCs in thin slices of cerebellar cortex (Figure 1B). To control for shared gradients of morphological variation in humans and mice^11,36^, we sampled PCs from the same region across species. Here, we chose Crus 1-2 lobules of the posterior mid-hemisphere because this region has been linked to multisensory integration and mixed selectivity or higher-order cognitive functions in mice and humans, respectively, and has selectively expanded in volume in primates^50^, suggesting that this region may be a key site for adaptation to species-specific functions. To capture broad evolutionary trends, we grouped species into clades for analysis: rodent (Mus), platyrrhine (Plat), cercopithecid (Cerco), hylobatid (Hylo), non-human hominid (NH-Hom), and human (Hsap) (n=3,3,5,4,9,3 specimens/clade).

**Table 1:** Attributes of the specimens used in this study.

| Species | Common Name | Sex | Age (yrs) | Rearing |
| --- | --- | --- | --- | --- |
| *Pan paniscus* | Bonobo | F | 31 | Hand |
| *Pan paniscus* | Bonobo | F | 28 |  |
| *Gorilla gorilla* | Gorilla | M | 44 | Hand |
| *Gorilla gorilla* | Gorilla | F | 31 | Hand |
| *Gorilla gorilla* | Gorilla | M | 46 | Hand |
| *Pongo pygmaeus* | Bornean orangutan | M | 40 | Wild |
| *Pongo abelii* | Sumatran orangutan | F | 50 | Captive |
| *Pan troglodytes* | Chimpanzee | M | 35 | Nursery |
| *Pan troglodytes* | Chimpanzee | F | 39 | Other - Captive Birth |
| *Symphalangus syndactylus* | Siamang | F | 27 |  |
| *Symphalangus syndactylus* | Siamang | M | 32 | Captive |
| *Nomascus leurogenys* | Northern white-cheeked gibbon | F | 45 | Wild |
| *Nomascus leurogenys* | Northern white-cheeked gibbon | F | 43 | Wild |
| *Macaca mulatta* | Rhesus macaque | M | 13 |  |
| *Macaca mulatta* | Rhesus macaque | M | 10 |  |
| *Macaca mulatta* | Rhesus macaque | F |  |  |
| *Papio anubis* | Olive baboon | F | 11 |  |
| *Papio anubis* | Olive baboon | M | 11 |  |
| *Callithrix jacchus* | Common marmoset | M | 3 |  |
| *Callithrix jacchus* | Common marmoset |  |  |  |
| *Callithrix geoffroyi* | White-faced marmoset | M | 8 |  |
| *Ateles geoffroyi* | Black-handed spider monkey | F | 25 |  |
| *Homo sapiens* | Human | F | 92 |  |
| *Homo sapiens* | Human | M | 47 |  |
| *Homo sapiens* | Human | F | 34 |  |

**Figure 1:**
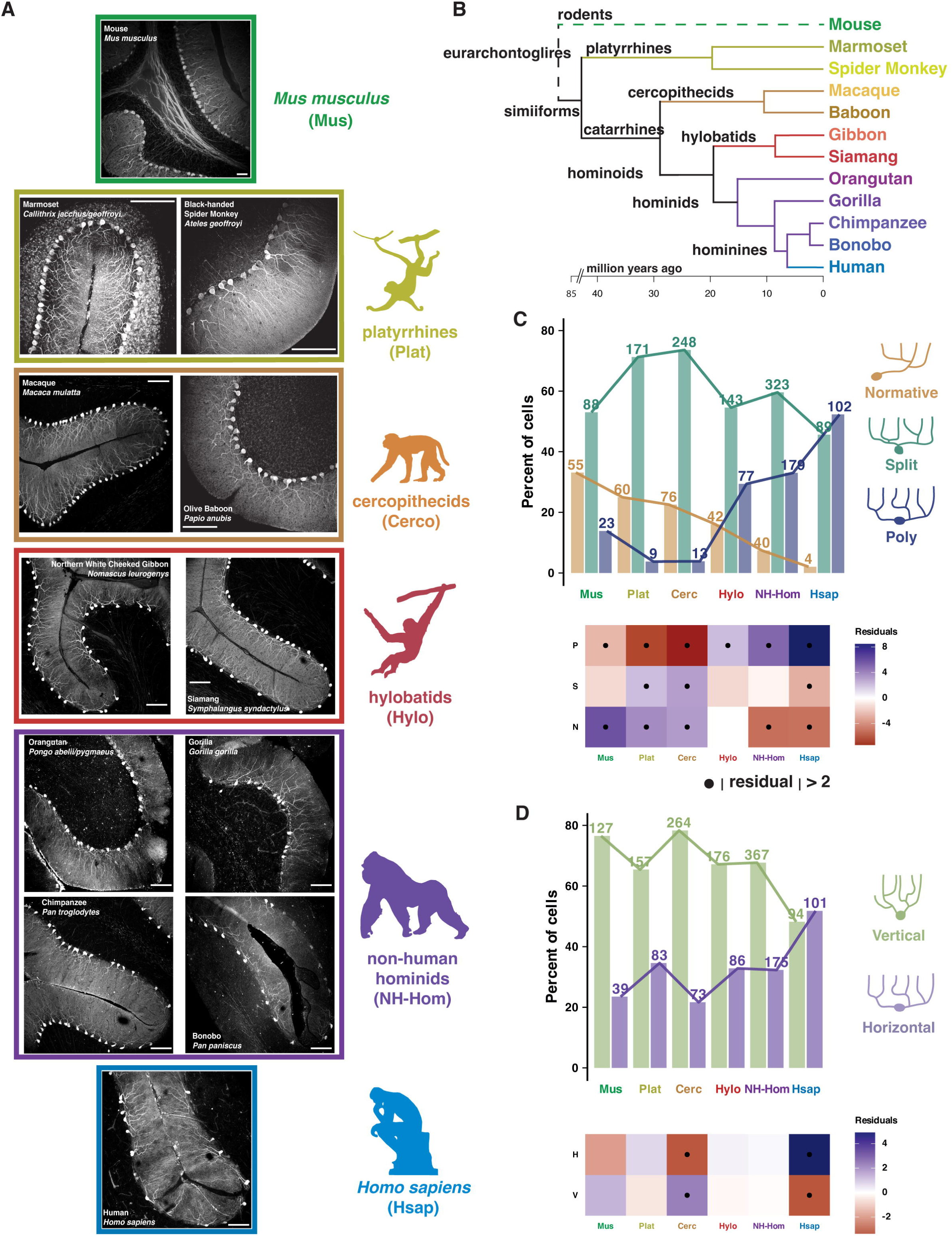
PC morphological subtype demographics shift across primate evolution. A) Sample confocal images from each species, organized by clade, of calbindin immunolabeled PCs. Note, total image scales varied somewhat with tissue size while resolution was maintained. Scale bars denote 200μm. B) Phylogenetic tree of the species sampled. C-D) Phylogenetic patterns in the distribution of PCs when categorized according to the type (Normative, Split, or Poly) and orientation (Vertical or Horizontal) of their primary dendrites using previously described criteria^11,36^. Clade order and connecting lines highlight the human evolutionary trajectory. Numbers of cells is noted above percentage bars. Heatmaps show χ² standard residuals with strong effects >|2| highlighted.

As primary dendrite morphology plays an important role in PC physiology^36,51,52^, we began by characterizing PCs according to our previous framework for defining primary dendrite structure: Normative (one primary dendrite), and two categories of multi-dendritic motif that are Split (one primary dendrite that bifurcates near the soma) or Poly (multiple primary dendrites emerging from the soma) (see Methods). Multi-dendritic (Split, Poly) morphologies are associated with CF multi-innervation and dendritic signal heterogeneity^11,36^, so these subtype categories serve to indicate the potential for complex CF-dependent signaling. Subtype distributions differed substantially across clades (Figure 1C; χ² = 323.7, p<0.001). While Crus 1-2 has the lowest expression of Normative PCs in mouse hemisphere^36^, mice nonetheless had more Normative PCs than any simiiform. The rate of Normative PCs drops across the human evolutionary trajectory, giving rise to increasing numbers of multi-dendritic PCs (Split or Poly). Primate clades exhibited distinct patterns in the expression of the multi-dendritic motif, where Split PCs strongly predominated in platyrrhine and cercopithecid monkeys, and Poly PCs exhibited stepwise jumps in frequency from monkeys (∼5%) to non-human apes (∼30%) and reached their highest prevalence in humans (∼55%). Species level patterns broadly mirrored these patterns but with some variation (Figure S1A).

We next examined dendritic orientation (Vertical versus Horizontal, χ² = 58, p<0.001) to capture geometric variation independent of primary dendrite number (Figure 1D). Horizontally oriented PCs were selectively enriched in humans (+6.3 standard residual, SR) and reduced in cercopithecids and mice (-4.5, -2.5 SR). Although orientation frequencies varied across species, humans were unique in having horizontal PCs represent the majority subtype (Figure S1B). PCs therefore become increasingly polydendritic and horizontally oriented along the human evolutionary trajectory (visualized in Figure 3), leading us to ask whether this categorical description masks further clade-specific changes in PC morphology.

### Human PC morphology is driven by a trend toward increased dendritic complexity and compartmentalization

We performed a parametric analysis of >1,700 PC morphologies across clades (160-540 cells across 3-9 brains per clade) and species (>100 cells and 2+ brains per species, except only one Black-Handed Spider Monkey). Because primary dendrites shape PC integration and are well preserved even in non-perfused archival tissue, we focused on quantifying geometric and branching properties of these larger dendrite structures. For each cell, we manually measured up to 9 morphological parameters—including trunk diameters, bifurcation distances, branching angles, and dendritic separation—for each primary dendrite (requiring replicate measurements for multi-dendritic cells) along with local cortical structure such as molecular and granule layer thickness and foliar location (Figure S2; Methods).

We first asked whether individual traits exhibit directional trends across primate phylogeny. The variation in several features converged on a common theme: increasing dendritic compartmentalization in the human trajectory. The distance from the soma to the first bifurcation (split distance) did not scale in proportion with increasing cortical thickness in monkeys and even shortened among apes (Figure S3A). When normalized to the surrounding tissue dimensions, therefore, split distance dropped progressively from mice to monkeys to apes and humans (Figure 2A). The primary bifurcation angle widened (Figure 2B). Together, these changes shift the divergence of a multi-dendritic structure to occur closer to the soma and expand the spatial separation of emerging compartments. This was reflected in the increasing number of primary dendrite compartments (Figure 2C), defined as the number of thick caliber dendrites originating through bifurcations close to the PC layer, and the horizontal distance between the centroids of the outer compartments (Figure 2D).

**Figure 2:**
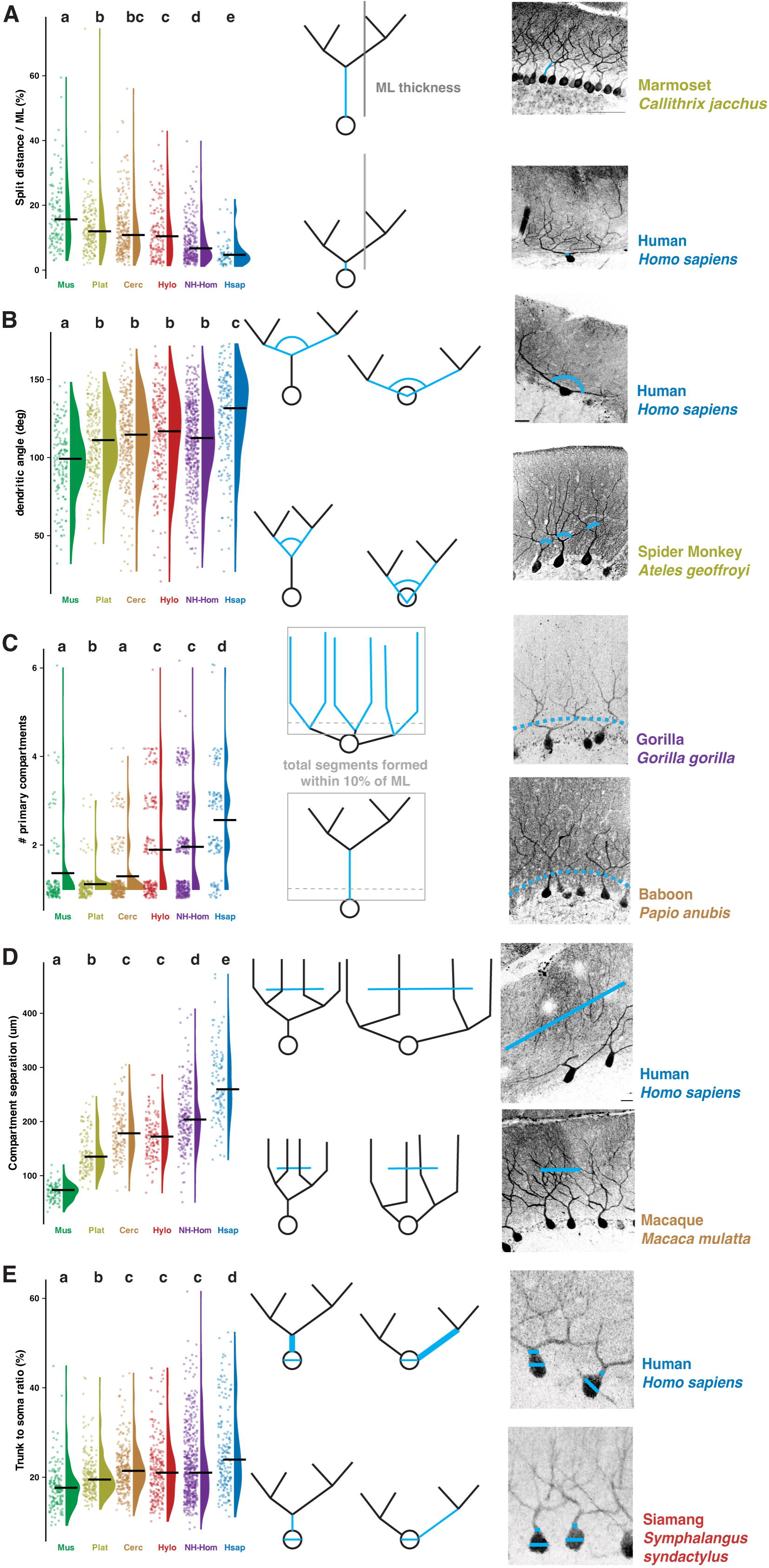
Human PCs diverge from other primate clades across morphological features. A) Distance to bifurcation normalized to total ML thickness of cells with one primary dendrite (here and below, letters indicate statistically distinct groups from a Dunn post-hoc test following pairwise Kruskal-Wallis). B) Angle of dendritic ramification from the bifurcation of a single primary dendrite or of multiple dendrites as they emerge from the somatic compartment. C) Number of primary dendrite compartments, defined by the number of dendrites emerging from the soma and bifurcating within the proximal 10% of the ML. Vertical and horizontal jitter is applied to visualize discrete values. D) Distance separating the centroids of the two outer dendritic compartments, measured at half the ML thickness. E) Ratio of somatic diameter to largest trunk thickness. Scale bar denotes 100μm.

Split PCs with short split distances had a broader range of trunk diameters (∼2-22μm) while those with long distances converged on a more conserved trunk diameter of ∼4-7μm across primates and including mice (Figure S3B). Consistent with this, human PCs, in which short splits predominate, had higher trunk to soma diameter ratios (Figure 2E). This supports the hypothesis that Normative, Split, and Poly PCs may lie on one developmental axis dictating the propensity for primary dendrites to rapidly diverge at a short distance from the soma, from a perisomatic dendritic extrusion, or even directly from the soma. An additional support for this hypothesis may be found in the presence of Poly PCs with large disparity between the thicknesses of their multiple primary trunks, possibly the result of an increasing drive to retain multiple nascent dendrites sprouting from the soma even if some are diminutive and would otherwise have retracted. This parameter exhibits a complex relationship across primate phylogeny (Figure S3K), whereby the trunk thickness disparity is large in non-hominoids (monkeys) with few Poly PCs, low in non-human hominoids (apes) where Poly PCs become common, and high again in humans where Poly PCs become the predominant subtype (Figure 1C). This developmental axis would control the formation of multiple primary dendrite compartments with differing signal mixing before the final integration at the somatic compartment.

Taken together, these individual trait variations produce a coherent shift in the morphologically average PC across primate evolution (Figure 3). For mono-dendritic (Normative and Split) PCs, this shift primarily represents a shortening split distance and the increasingly proximal emergence of additional primary compartments. In the case of multi-dendritic (Poly) PCs, the notable morphological shift is an increasingly obtuse dendritic angle as dendrites project more horizontally, rendering primary dendritic compartments more separable.

**Figure 3:**
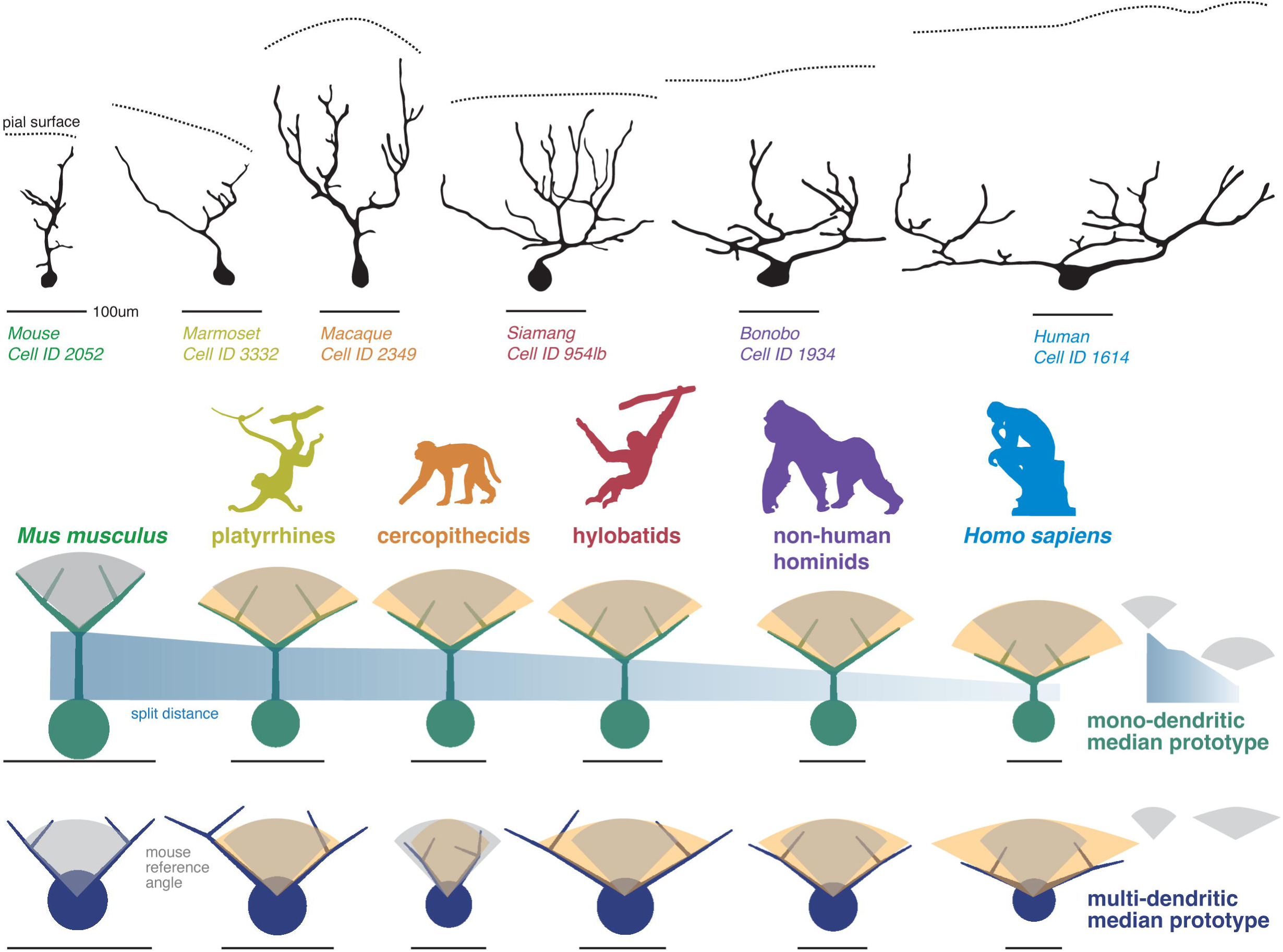
Morphological shifts along the human evolutionary trajectory. Manually traced primary dendrites of representative PCs for each clade (all cells on the same scale). Cells were chosen from the broader dataset as those with features closest to clade medians. Below are schematized ‘prototype’ models of the primary dendrite architecture for mono-dendritic (Normative and Split) and multi-dendritic (Poly) PCs using the median trait values. Cell scales are normalized to their respective median ML thickness to indicate the relative size of each morphological feature in the cortex. For example, increasing soma size is often sublinear to the expanding ML thickness, producing relatively smaller somata in humans than mice. Scale bars denote 50μm.

Although directional shifts were evident, many morphological traits displayed substantial cell-to-cell variability within clades (Figure S3) and species (Figure S4-5), challenging the notion that PC morphology is highly stereotyped. We analyzed the intrinsic variation of each morphological feature to determine whether it is tightly or permissively regulated. To appropriately compare variables across scales and units, we quantified intrinsic dispersion using dimensionless metrics: robust coefficient of variation, Shannon entropy (for discrete traits), fraction of total observed range occupied, and normalized standard deviation (Methods; Figure S6). Several traits exhibited high dispersion—including split distance, poly-dendrite angle, trunk disparity, and secondary bifurcation distance—suggesting permissive regulation. In contrast, soma diameter, secondary angle, and secondary bifurcation angle were tightly constrained across phylogeny. Notably, many parameters shifted in regulatory permissiveness along the evolutionary trajectory, suggesting that human evolution extended beyond directional shifts in median values into expanded morphological variability in part resulting from the greater degrees of freedom provided by increasing compartment number. Thus, human PC evolution entails not only a structural shift relative to other simiiforms but expansion of the morphological configurations available.

### PC morphology lies on a high dimensional morphospace

The high variability across many traits raised the possibility that clade-specific adaptation may act upon combinations of morphological features rather than individual parameters. To test this, we analyzed PC morphology in multivariate morphospace defined by each cell’s profile across all measured traits (up to 17 features/cell) and excluding cell identifiers (e.g. species, clade, or assigned morphological category). Pairwise distances between cells were calculated in morphospace and visualized using principal coordinate analysis (PCoA). Although some structure was evident in the first few PCoA axes, modest clustering emerged only along the first axis, which primarily reflected the presence or absence of multiple primary dendrites (Poly vs non-Poly; Figure 4A). All other morphological features exhibited weak associations with early axes (e.g. Figure 4B), and each axis explained only a small fraction of variance, with only the first 3-4 axes exceeding null shuffled datasets (Figure 4C). This pattern is characteristic of high-dimensional systems in which variables do not strongly covary (Figure 4D) and instead represent largely independent axes of variation. Thus, PC morphology occupies a high-dimensional morphospace consistent with independent developmental control over individual morphological features.

**Figure 4:**
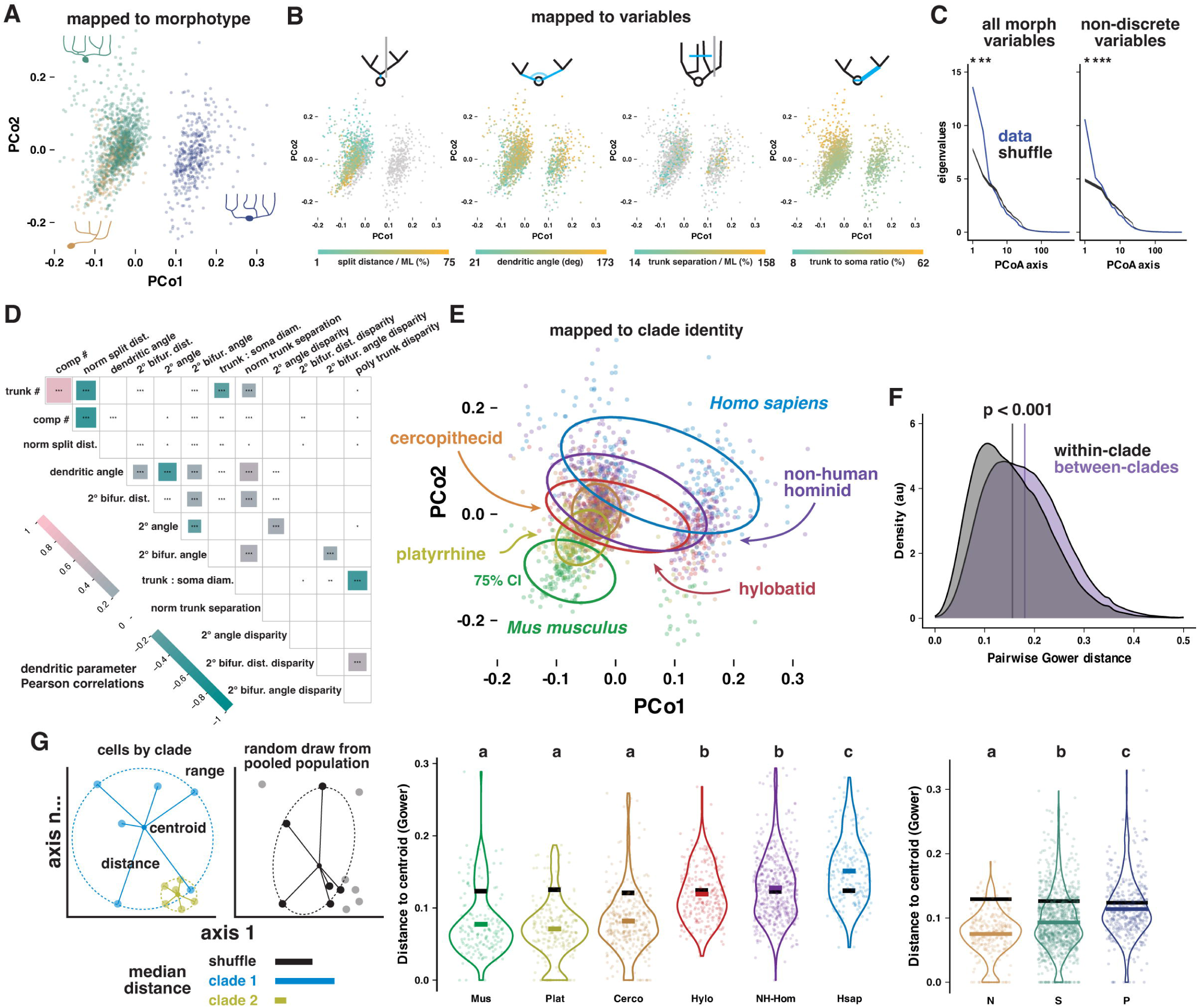
PC morphology is high dimensional and varies across primate phylogeny. A) Visualization of PC positions on the first two axes of morphospace following dimensionality reduction by PCoA colored by reattributed morphological category. The first axis largely encodes whether or not a PC is Polydendritic. B) Examples of individual morphological traits being poorly predicted by the first two PCoAs. C) PC morphospace is high dimensional; early axes capture minimal variance and otherwise resemble a shuffled dataset. D) Matrix of inter-variable Pearson correlations showing minimal covariance of morphological traits. Values below ±0.2 removed for clarity. E) As in A, but colored according to reattributed clade identity, illustrating PCs from the same clade tend to occupy similar regions of morphospace. F) Pairwise PC distances in morphospace for pairs in the same or different clades. G) Simplified schematic of the dispersion analysis (left) and distributions of distances of PCs from their respective clade (center) or morphology (right) centroids. Letters indicate statistically distinct groups from a Dunn post-hoc test following pairwise Kruskal-Wallis.

Despite independent variance across traits, reattributing clade identity to all cells revealed morphospace occupancy shifts across phylogeny (Figure 4E). We compared pairwise distances between cells within or between clades to show that PCs from the same clade occupied more similar regions of morphospace than PCs from different clades (0.156 vs 0.181, p < 0.001, Wilcoxon rank sum test; Figure 4F). Clade identity may therefore be reflected in multivariate shifts in morphology, which was supported by the dispersion of cells around their clade’s morphospace centroid (Figure 4G, left schematic). Dispersion was low in mice and platyrrhine and cercopithecid monkeys, indicating relatively stereotyped morphologies (Figure 4G). However, dispersion increased progressively in non-human hominids and ultimately humans, whose PCs spanned the broadest range of morphospace. This pattern was recapitulated at the species level (Figure S7A). Greater morphospace dispersion may arise from increased expression of specific morphological subtypes (Figure 1C-D). Consistent with this, Poly PCs occupied a broader region of morphospace than Normative or Split PCs (Figure 4H) as did horizontally oriented PCs compared with vertical PCs (Figure S7B). Both features increase the geometric degrees of freedom available to the broader dendritic architecture. These findings suggest that primate evolution has favored a shifted and expanded range of morphological configurations through increased dendritic area and compartmentalization rather than uniform shifts in a single structural parameter.

### Identifying nodes of evolutionary adaptation across primates

We next asked whether primate clades occupy distinguishable regions of morphospace. If evolutionary divergence altered PC morphology systematically, then morphospace profiles alone should predict a cell’s clade of origin. To test this, we trained a random forest classifier using all morphometric parameters and clade labels. Given the relatively low sample size after randomly downsampling each clade to the least number across clades (166 cells), we used 5-fold cross-validation (i.e. iteratively switching which 20% of the data was held out of training for testing so the full dataset was tested). The resulting model predicted clade identity above chance for all groups (Figure 5A; chance is 17%) with the highest accuracy for mouse (67%) and human PCs (61%), followed by platyrrhine (46%) and cercopithecid (42%) monkeys. Predictions were weakest for non-human hominids (31%). Hylobatids were predicted at chance level (18%), perhaps occupying an intermediate position and lacking distinctive traits. On the other hand, the low confusion among other clades indicates divergence through the acquisition of distinguishing morphological features rather than modest shifts in median trait values alone. Classification accuracy declined at the species level with more groups and less data per group, but similar structure persisted (Figure S7C–D).

**Figure 5:**
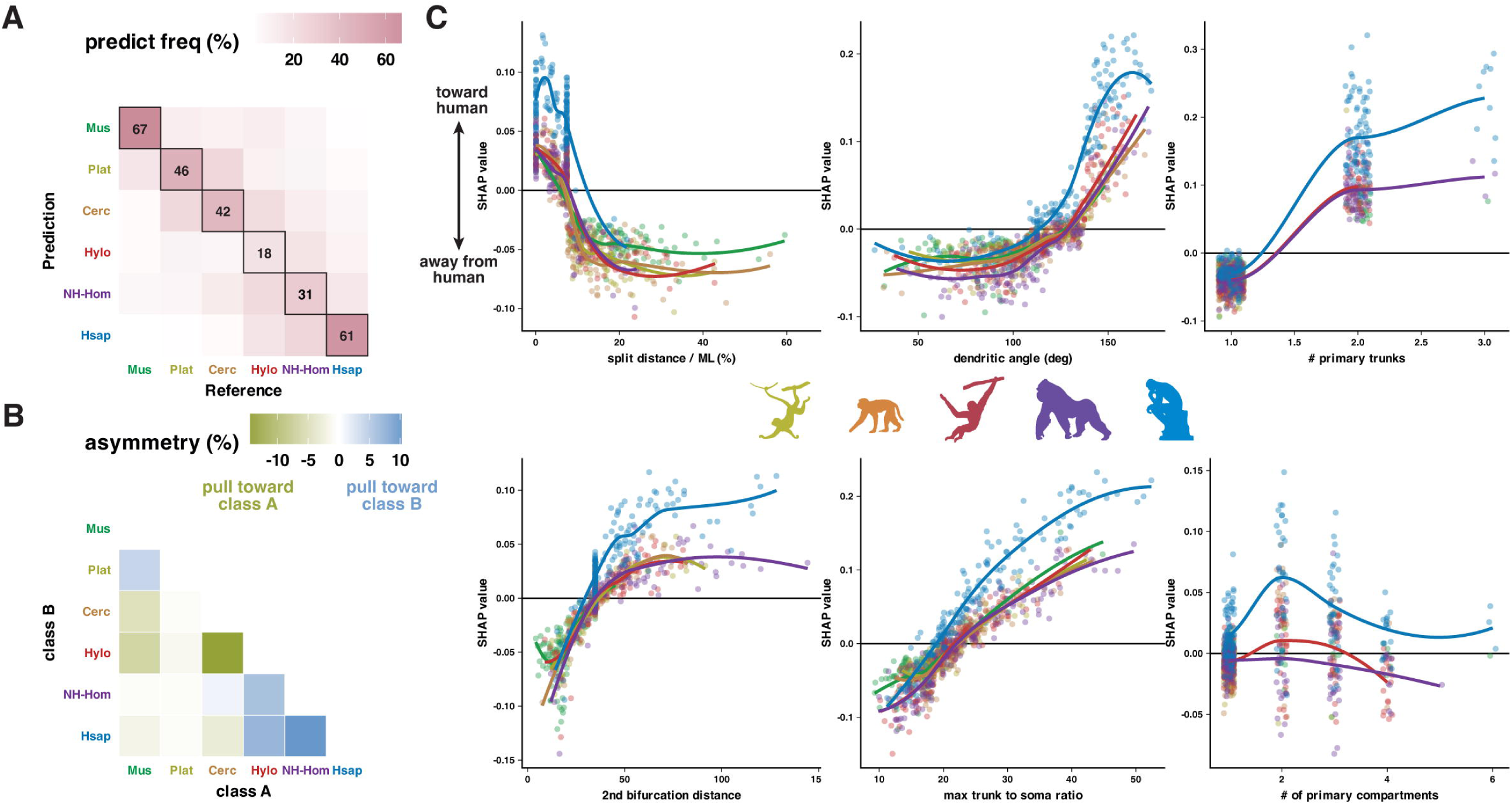
Morphological features predict primate clade identity. A) Random forest classifier predicting clade identity from PC morphology. Prediction accuracy exceeds chance (17%) for all clades, with the greatest separability for mice, humans, and platyrrhines and the lowest for hylobatids. B) Characterization of the off-diagonal prediction asymmetry in (A), measured as the difference between the frequency of clade A cells predicted to be clade B and the opposite. Asymmetry reveals directional relationships between clades, where negative values (olive) indicate clade pairs where prediction tends to pull toward class A. C) Shapley Additive Explanations (SHAP) analysis identifies the morphological features most strongly driving the classification of each cell as either belonging to a target class (clade) or not, here the target class was set to human. Positive and negative values are assigned to variables that either increased or decreased the probability that the cell was predicted to be human. The size of the value quantifies how strongly the variable drove the final prediction. Comparing the raw measurement and SHAP values of the strongest predictor variables across cells reveals the aspects of each variable (e.g. dendritic angles > 125, split distance ratios < 10%) that most strongly impact the prediction that a cell is human. By indicating true cell identities, these figures further demonstrate the specific dendritic features that are most uniquely characteristic of human PCs and therefore drive the greatest separation between predictions for human vs non-human cells. The same analysis for non-human clades is shown in Figure S8.

Inaccurate predictions (off-diagonal values) were also informative as they indicated which clades overlap most in morphospace. Mouse PCs were most often confused with platyrrhine monkeys and both monkey clades separated from apes. These cross-clade misclassifications were sometimes asymmetric (Figure 5B), demonstrating some directionality in the relationship that can indicate their relative positioning in morphospace. For example, primate PCs are more often predicted to be cercopithecid PCs than cercopithecid PCs are predicted to be any other clade (negative asymmetry for Cerc → Hylo, NH-Hom, Hsap and positive asymmetry for Plat → Cerc). This suggests that cercopithecid PCs, which are relatively stereotypical (low centroid dispersion), may represent a compact origin in morphospace from which other primate clades broadened their occupancy of morphospace while retaining overlap with cercopithecids. As platyrrhines are similarly compact but more distinct (lower confusion that is more reciprocal), this may be consistent with platyrrhines retaining more ancestral features while cercopithecids evolved more rapidly before diverging from hominoids.

We next decomposed the morphological features that most informed the classifier’s assignment using Shapley additive explanations analysis (SHAP; Methods) to generate a metric of each variable’s role in driving target class choice. Comparing the variable SHAP metric with each variable’s raw values highlights the aspects of each feature that pushed the prediction either toward or away from a particular class. For example, PCs were increasingly predicted to be human if they had shorter split distances, wider primary dendrite angles, longer distances to the secondary bifurcation, and increased thickness of the primary dendrite compared with the somatic size (Figure 5C). We performed the same analysis across clades to uncover the morphological characteristics that most distinguished each clade (Figure S8). Together, these results demonstrate that primate clades occupy partially separable regions of morphospace defined by specific morphological axes, revealing the traits that represent the strongest nodes for lineage-specific evolutionary divergence.

### Morphological features are not constrained by tissue foliation or allometric factors

We next evaluated whether PC morphology is shaped by mechanical constraints imposed by foliation or allometric scale^3,4^. PC dendrites may ramify in directions congruent with the mechanical effects of cortical foliation^9,53^, but it is not clear whether this would predict horizontal stretching of PC dendrites in the gyrus or the sulcus. We classified whether each cell resided along a gyral lip where the Granule Cell layer (GCL) pinches relative to the Molecular layer (ML), in the depth of a sulcus where the GCL stretches relative to the ML, or along a bank region where laminar contortion relaxes (Figure 6A). The prevalence of Normative, Split, or Poly PCs varied somewhat by foliar location across clades (Figure S9D), in keeping with previous findings in mice and humans^11,36^. PC orientation exhibited more robust region specificity, with horizontal structures were favored in the sulcus across clades (Figure 6A), possibly indicating a stronger response to the thinning and stretching of the GCL. Echoing these findings, only dendritic angle varied according to foliar region (Figure 6B), though clade-level differences nonetheless exist in matching foliar sub-regions (Figure S9E). Together, these analyses indicate that neither cortical scale nor foliar architecture sufficiently explain the phylogenetic morphological divergence we observed, supporting the hypothesis that PC morphological adaptation may be driven more by changing functional demands rather than mechanical constraints.

**Figure 6:**
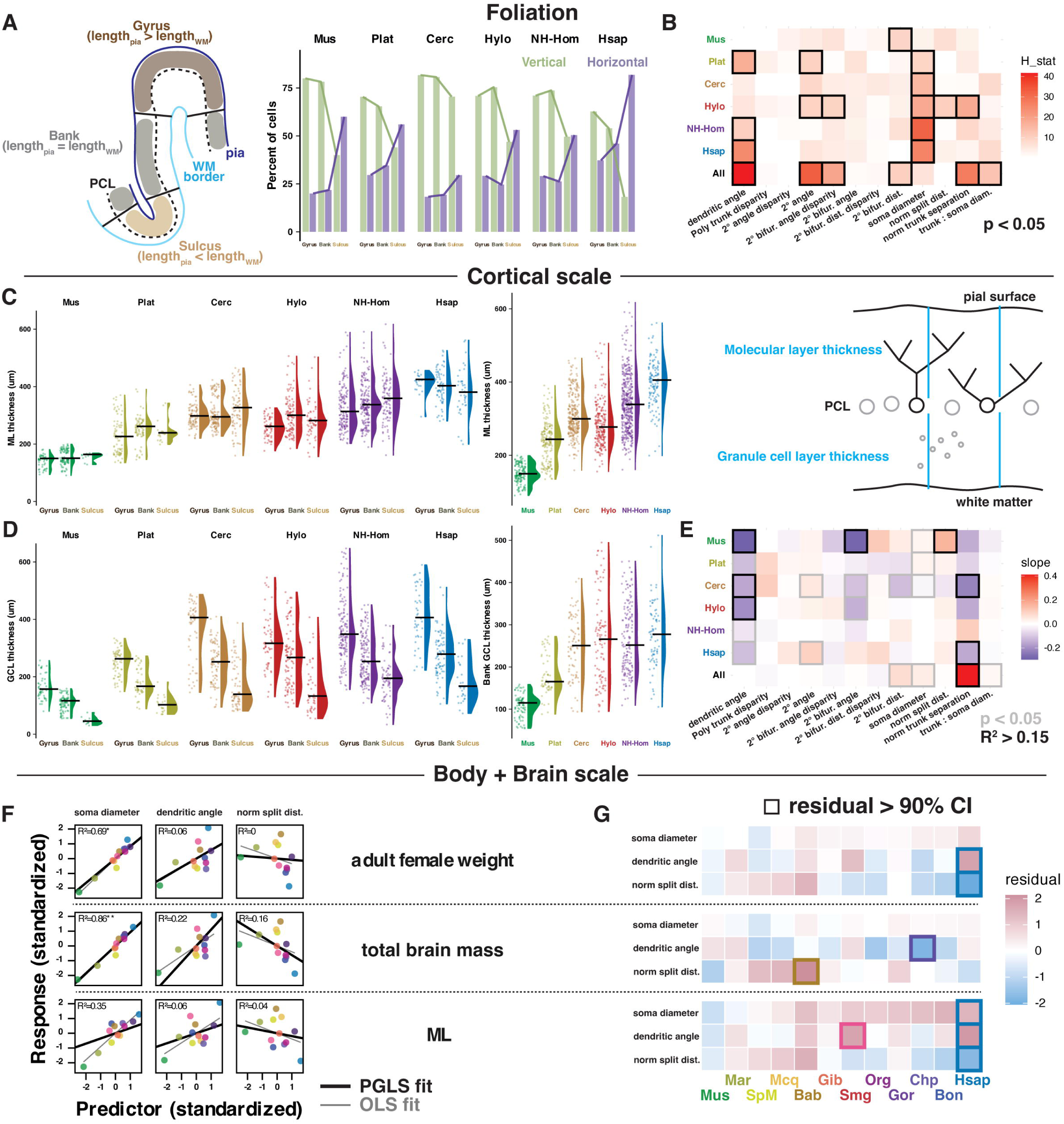
Phylogenetic variation in PC morphology is broadly foliation invariant and not allometrically scaled. A) Schematized foliar definitions (left) and distribution of PC categories by clade and foliar region. B) Heatmap table highlighting H-stat of pairwise Kruskal-Wallis tests of difference for each morphological variable by the foliar region within each clade and as a total population. Black outline denotes a significant relationship. C-D) Local ML and GCL thickness measured at each cell by clade and foliar region. E) Heatmap table highlighting the slope of the linear regression of each variable to ML thickness within each clade and as a total population. Grey outline denotes a significant relationship, and black outline denotes cases where the linear model explains more than 15% of the variance. F) Example morphological variables regressed against allometric factors of body mass, brain mass, and ML thickness with phylogenetic generalized least squares (PGLS, black) or ordinary least squares (OLS, grey) models. Points are the median values for each species. Body and brain size predictors were log scaled. For comparison, all predictors and response variables were standardized (centered and divided by the population standard deviation). G) Species residuals from PGLS fits. Borders denote residuals exceeding the 90^th^ percentile of null data simulating Brownian motion.

As previously known, total brain and cerebellum mass scale with body size in accordance with an allometric power law across primates, with the notable exception of humans (Figure S10A). For each clade, we included smaller members (e.g. marmoset, macaque, and gibbon) and larger members (e.g. spider monkey, baboon, and siamang). As the cerebellar cortex is the most direct constraint on PC size, we assessed whether morphology within and across species was predicted by cortical thickness or foliation. Where permitted by tissue quality, we measured the thickness of the ML where PC dendrites reside and the GCL where the main dendritic input population resides and could conceivably influence PC morphology. Median ML thickness, and to a lesser extent GCL thickness, scaled with cerebellar mass (Figure S10A), but was invariant to foliar location (Figure 6C) while the GCL thinned in the sulcus and expanded in the gyrus (Figure 6D). ML thickness did not predict position in morphospace (PCoA axis 2 R² < 0.25, all other axes R² < 0.1; Figure S9A) or individual measurements (Figure 6E; Figure S10A-B). Trunk separation had a modest positive correlation with ML across clades (Figure S9B), but within clades slopes were weak or reversed (Figure S9C), illustrating a Simpson’s paradox whereby an across-group trend obscures an opposite within-group trend. Thus, population-level tissue scaling does not play a role in the phylogenetic trends we observe in PC morphology.

Finally, we asked whether PC morphology reflects allometric scaling with body or brain mass. These allometric factors were poor predictors of most morphological features, even accounting for phylogenetic covariance (Figure 6F, S10A-B). Phylogeny alone was also a poor predictor, with many variables showing stronger conservation than predicted by Brownian motion (Blomberg’s K > 1.5) or little detectable phylogenetic signal (K < 0.5) while only some show phylogenetic signal (Figure S10C). Intriguingly, brain mass often provided the best model fit to PC morphology, which was otherwise largely invariant to body mass or local cortical thickness. Human PCs deviated most strongly from the phylogenetically controlled regression to body and cortex size, but their morphology was often well aligned to brain mass (Figure 6G, S10D).

## Discussion

Adaptive remodeling has defined the central nervous system since its ancient bilaterian origin. Although speciation can drive the remodeling of neural circuits supporting lineage-specific behavior, regions dedicated to conserved sensory transformations or internal state regulation frequently preserve their core architecture. For example, prolonged telencephalic development in mammals promoted expansions in lamination and circuit complexity, factors associated with more integrative and flexible capacities for decision-making and learning^54,55^. On the other hand, the mesencephalic colliculus retains its core anatomical and functional arrangement with some variation in input weight and lamination^56^. Developmental patterning of the hypothalamus relies on gene regulatory programs traceable to basal chordates such as *Ciona* sea squirts^57^. Considering the factors governing change and preservation in the brain—geometrical complexity and shifting functional demands—the cerebellum presents a surprising case where form and function are thought to be strongly conserved across vertebrates since their emergence ∼530million years ago. The cerebellum has a complex cortical circuit that directly subserves behaviors as numerous and diverse as the species performing them: corollary charge subtraction in electric fish proprioception^58^; vocalization precision and learning in songbirds^59^; undulating, rectilinear, or legged movement among lizards and snakes (squamates)^60,61^; language and higher-order cognitive performance in humans^48,62^; and further specializations that currently evade investigation even among our closest relatives, ranging from specialized arboreal locomotion techniques, such as brachiation in gibbons and prehensile tail use in spider monkeys, to poly-dexterous limb use among non-human great apes.

At what levels of organization does adaptation arise in the cerebellum? Reports of anatomical specialization largely involve gross morphology such as foliation (largely consistent in eutherians^63^ and birds, but absent in squamates^3,61^ and other vertebrates), scale (volume increases matching neocortical expansion^64,65^ and duplicated archetypal subnuclei expanding deep cerebellar nuclei output channels^66^), or gene-expression profile variation (increased PC abundance in early human fetal growth compared with rodents and marsupials^33,67^, region-specific transcriptional divergence tracking primate phylogenetic relationships^68^). Cellular morphology and physiology, however, is considered highly stereotyped within and across species^69,70^. Yet, recent studies show that PC morphology varies regionally^11,36,38^ and between humans and mice^11,36,37^, suggesting that PC morphology may be more permissively regulated than previously appreciated. PCs could therefore act as a node for cerebellar adaptation amid diversifying behavioral and sensorimotor demands.

Here we build on our recent framework for physiologically relevant categorization of PC morphology with a parametric characterization of PC primary dendrite morphology across primates. We found that simiiform primate evolution drove expanded dendritic compartmentalization along the human lineage, expanding the morphological degrees of freedom, through multiple factors that may be independently regulated in development. Evolutionary adaptation of primate PCs therefore appears to have proceeded not by shifting a single morphological axis but by expanding the dimensionality of allowable structural, and perhaps functional, configurations. Simiiforms represent a narrow segment of cerebellar evolution, suggesting that the cerebellar cortex is more anatomically diverse and rapidly evolving than previously appreciated.

### PC morphology as a regulator of circuit physiology

Our findings bear directly on theories of cerebellar computation, which generally take as a premise that the cerebellar circuit is uniform and generates a generalizable algorithm^8,71–74^ or that differing properties of input to the uniform circuit can produce some algorithmic variation^75–77^. We previously linked a multi-dendritic motif to CF multi-innervation and heterogeneous dendritic integration in mice^36^, with emerging evidence for a conserved structure-function relationship in humans, where this motif is abundantly expressed^11^. The geometrically precise cerebellar circuit architecture provides a structured substrate in which small modifications in dendritic morphology can have significant ramifications.

From a geometric perspective, the horizontal ramification of PC dendrites could have implications for several circuit functions. GCs receive clustered mossy fiber inputs^78^, producing spatio-functional clustering of the PFs that pass orthogonally through the PC arbor^79^. Horizontally separated dendritic compartments would then allow a multi-dendritic PC to survey distinct PF receptive fields (RFs). As PC intrinsic excitability gates the spatial distribution of PF synaptic plasticity^80–82^, dendritic arrangement may serve a critical role in compartmentalizing PF signal integration^52^ particularly in the case of morphologies producing spatially segregated compartments. Variable trunk thickness or bifurcation distance may regulate the dendritic integration of PF RFs and the contiguity of dendritic integration with the somatic compartment where signals are ultimately mixed to generate output. PF-LTD-dependent cerebellar learning largely relies on PF-CF coincidence^83^. The critical prerequisite for accurate plasticity is, therefore, matching RFs between CFs and PFs^80,83^ to prevent coincident PF weights from being adjusted inappropriately. PCs with broad, compartmentalized dendritic arbors and distinct PF RFs may require multiple CFs to match RFs between contextual PF and supervisory CF signals.

We do not know how CF multi-innervation impacts the cortical circuit, the olivocorticonuclear loop, or extra-cerebellar function. Cerebellar supervised learning by local CF signaling is conveyed to somatosensory cortex where it provides instructive signals for pyramidal neuron plasticity via specific effects on interneuron subtypes^84^. The observed variety in CF-PC input strength and numerosity resulting from normal development^36,85,86^ appears regulated to a certain range as excess CF-PC input strength and number have been linked to syndromic autism-like traits and spinocerebellar ataxia in mice^44,45,87,88^ and essential tremor in humans^89^. Systematic alterations in PC morphology are also linked to cerebellar diseases in these species^44,45,87,88,90–93^.

### PC morphology as a key node for behavioral adaptation during cerebellar evolution

Across vertebrate evolution, neural systems meet changing behavioral demands by balancing architectural conservation with functional adaptation. Our findings indicate that the cerebellum achieves this balance by retaining most of its geometrically precise circuit scaffolding while permitting flexibility at the level of its principal neuron morphology, where small structural deviations may confer outsized functional variation at implementation and algorithmic levels. PC primary dendrite structure therefore represents a node at which evolutionary pressure may tune computation without disrupting global circuit organization.

The mechanisms driving these morphological shifts remain unknown. Candidate mechanisms include cell-autonomous molecular pathways such as: self-avoidance factors like clustered protocadherins^94,95^, upstream transcription factor CTCF ^96^, *RORa* or Slit2/Robo2 ^97^; axon and dendrite polarity regulators (Liver Kinase B1 acting on Slit2)^98^; or atypical proteinase C (aPKC) control of golgi apparatus localization^99,100^. The cerebellum undergoes prolonged postnatal development, for which human-specific adaptations to regulate metabolism and synapse formation were recently described^68,101^. Yearslong postnatal refinement may render PC dendritic elaboration susceptible to activity-dependent mechanisms involving PF input^102^, intrinsic PC activity^103^, or a reciprocal stabilization and retention of multiple nascent dendrites by multiple competing CFs. Collectively, our findings echo a broader principle that evolutionary innovation in neural systems often arises not through wholesale remodeling, but through modification of key circuit elements.

---

**Figure S1:**
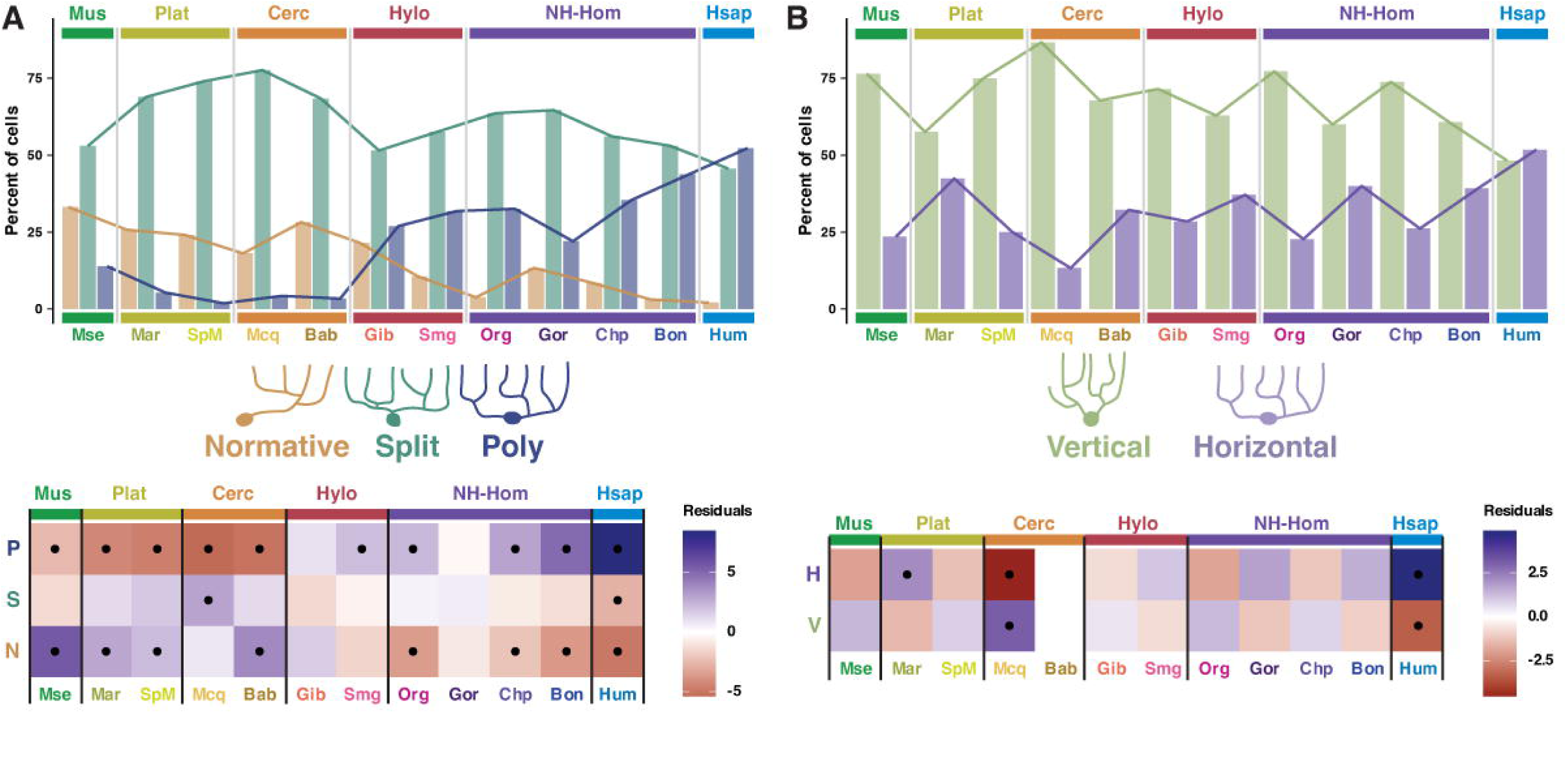
PC morphological categories vary across clade and species. A-B) Heatmaps of the standardized Pearson’s residuals (below the pooled median expectation in red, above expectation in blue) from chi-squared tests for differences in the prevalence of morphological (A) and orientation (B) subtypes across primate clades. Here and below, dots indicate strong effects where the residuals exceed ±2. C-D) Morphological category and orientation subtype distributions (top) and standardized residuals (bottom) across all species tested.

**Figure S2:**
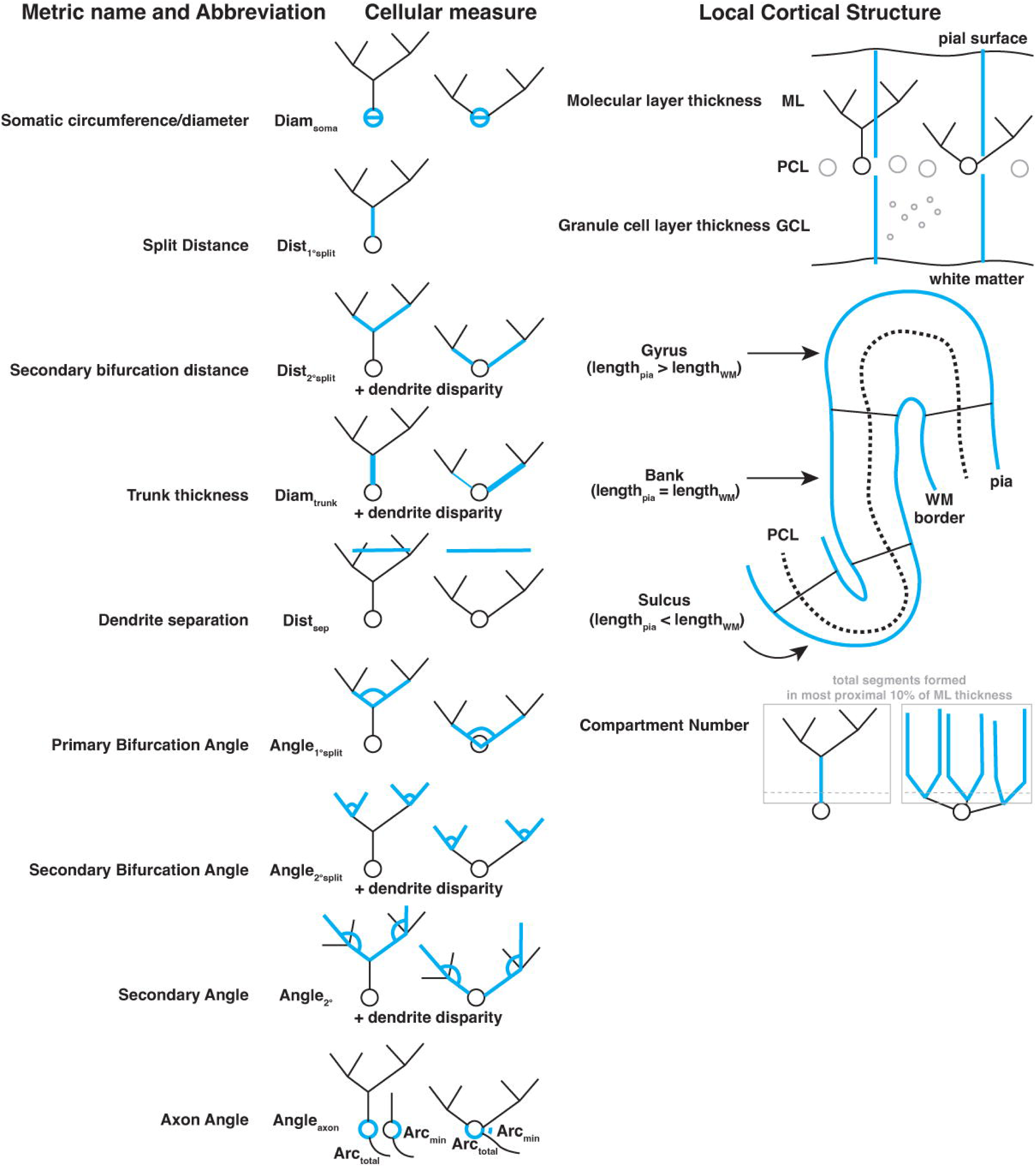
Outline of measured parameters describing morphology and local cortical structure.

**Figure S3:**
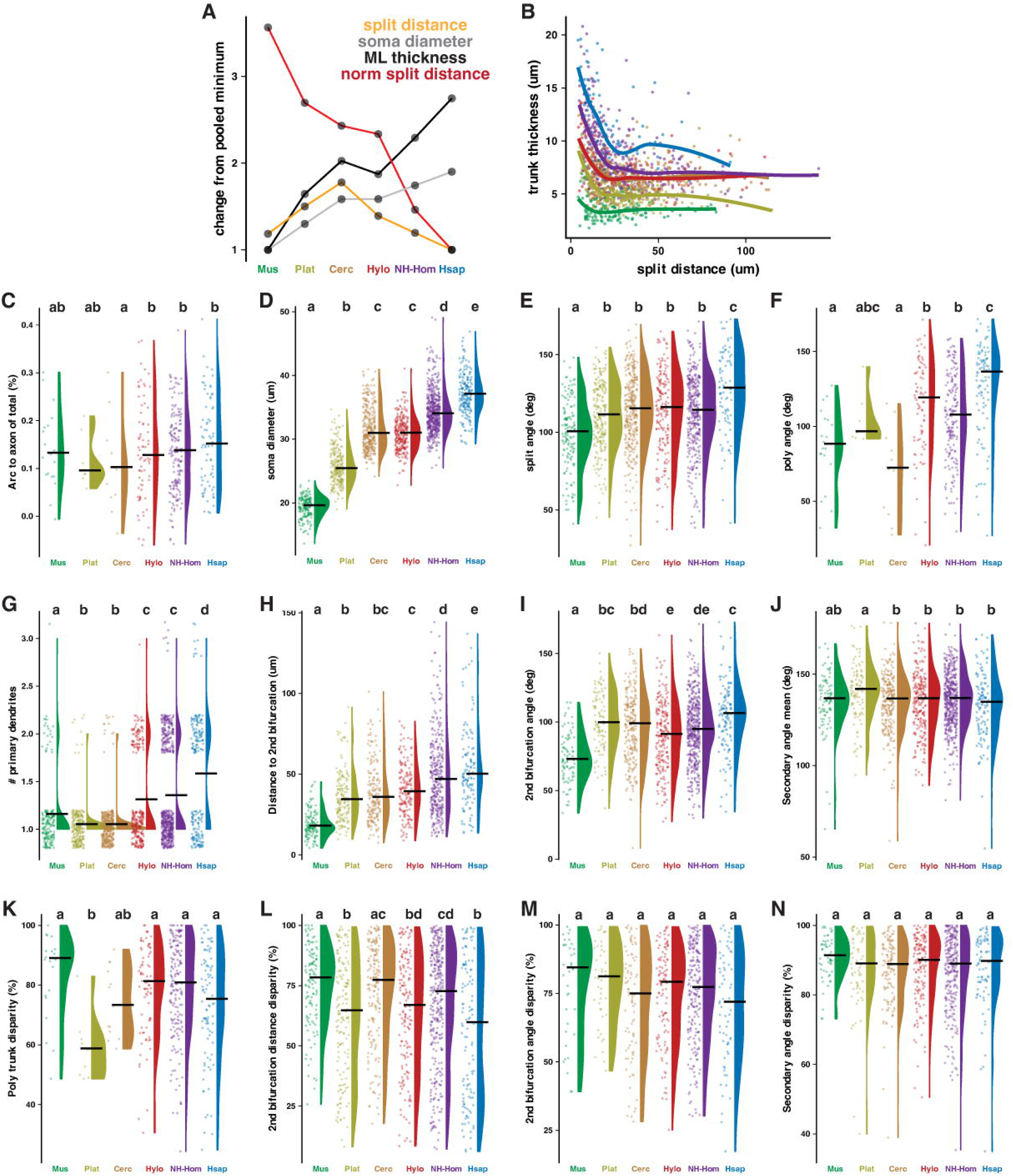
Characterizing morphological change by phylogenetic distance. A) Clade variation in split distance, somatic diameter, and ML thickness highlighting lack of proportional scaling of split distance and divergence of split distances normalized to ML thickness. B) Trunk thickness as a function of distance to the first bifurcation for PCs with one primary dendrite. Typical trunk thicknesses range by ∼7.5μm at low split distances (9-16.5μm) but converge on a narrower range (5-7μm) for PCs with longer split distances. C-N) Raw values by clade for additional variables.

**Figure S4:**
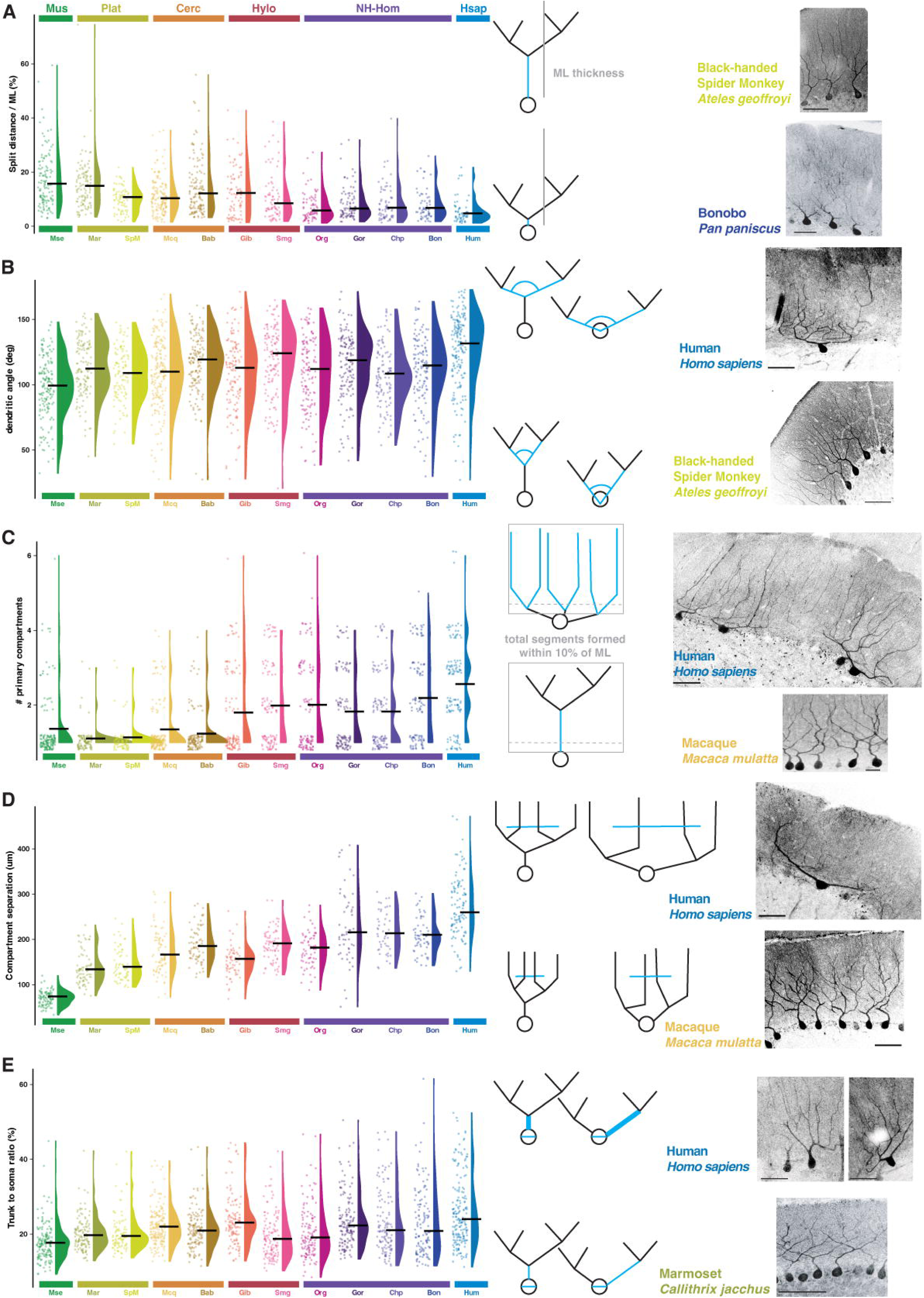
Morphological features showing the most consistent trends across species. Raw values as in Figure 2 but separated by individual species. A) Distance to bifurcation normalized to total ML thickness of cells with one primary dendrite. B) Angle of dendritic ramification from the bifurcation of a single primary dendrite or of multiple dendrites as they emerge from the somatic compartment. C) Number of primary dendrite compartments, defined by the number of dendrites emerging from the soma and bifurcating within the proximal 10% of the ML. Vertical and horizontal jitter is applied to visualize discrete values. D) Distance separation the centroids of the two outer dendritic compartments. E) Ratio of somatic diameter to largest trunk thickness.

**Figure S5:**
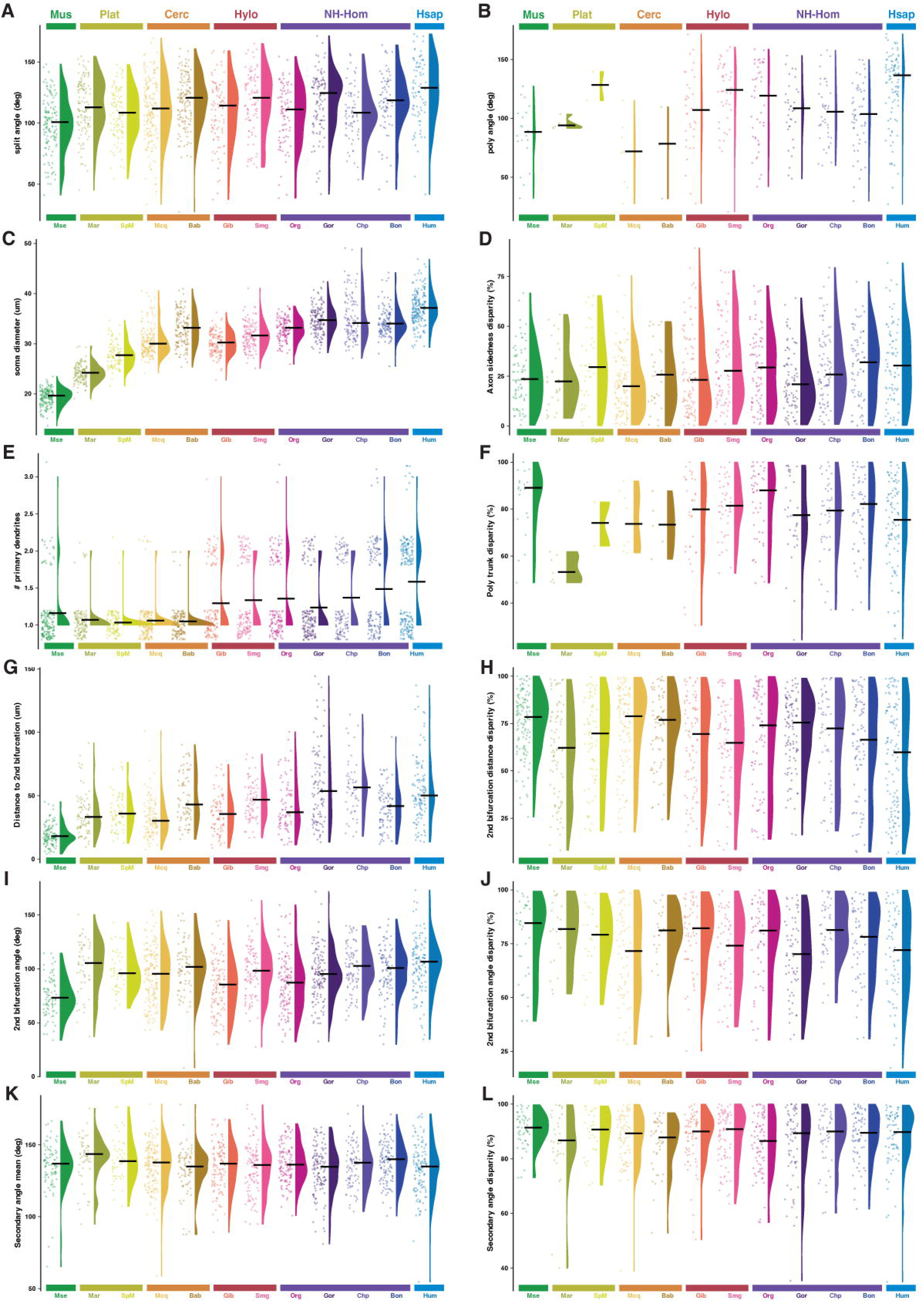
Additional morphological features across species. Raw values as in Figure S3D-O of additional morphological variables separated by individual species.

**Figure S6:**
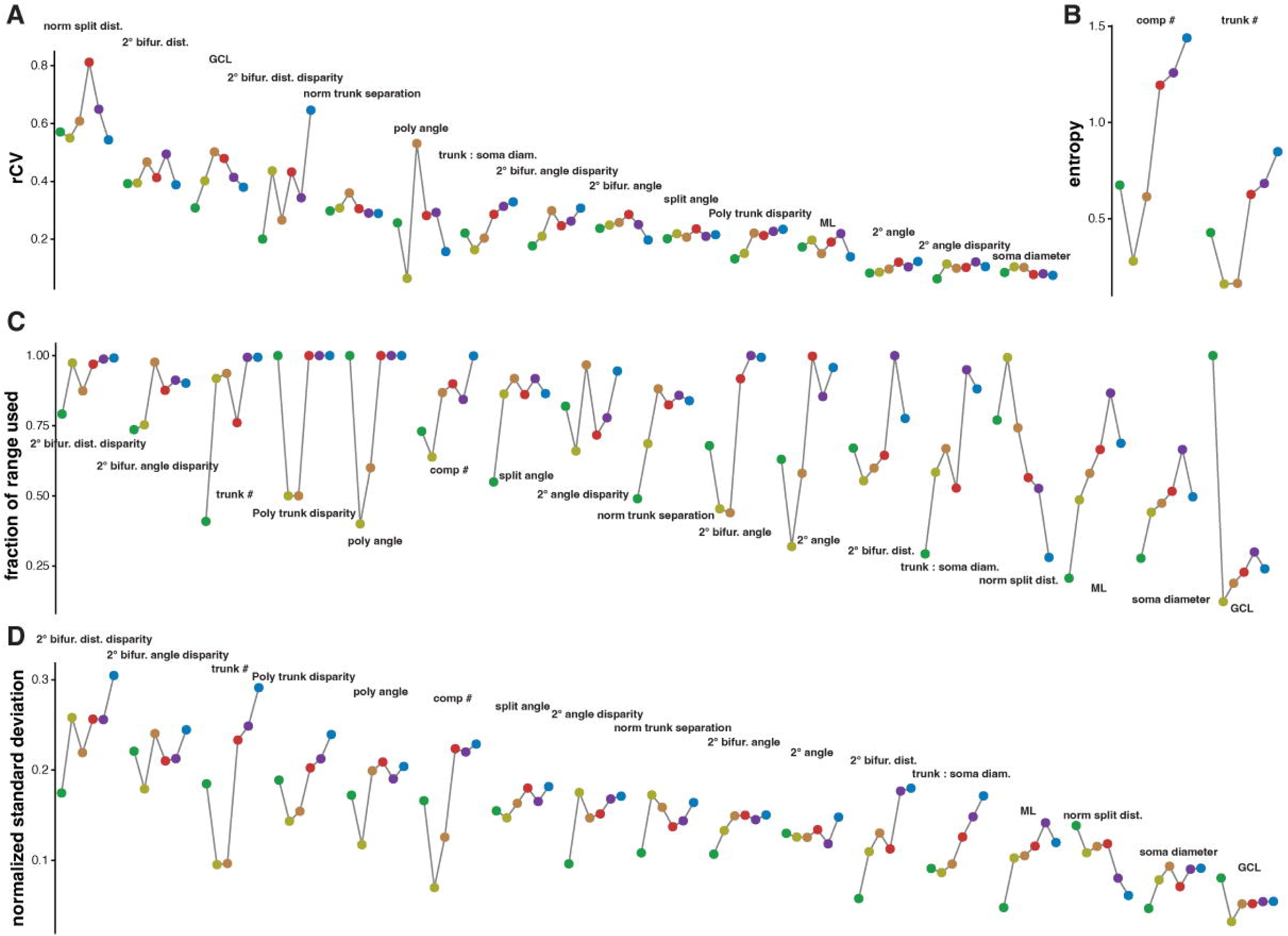
Assessing morphological variables with tight or permissive regulation within and across clades. A) Robust coefficient of variation (calculated on the normalized medians) of each morphological parameter across PCs in each primate clade. Sorted left to right by descending mean rCV across clades. B) Shannon entropy of discrete variables. C) As in (A), for the calculated fraction of the total observed range across all PCs that is occupied by PCs in each clade. D) As in (A), for the calculated normalized standard deviation.

**Figure S7:**
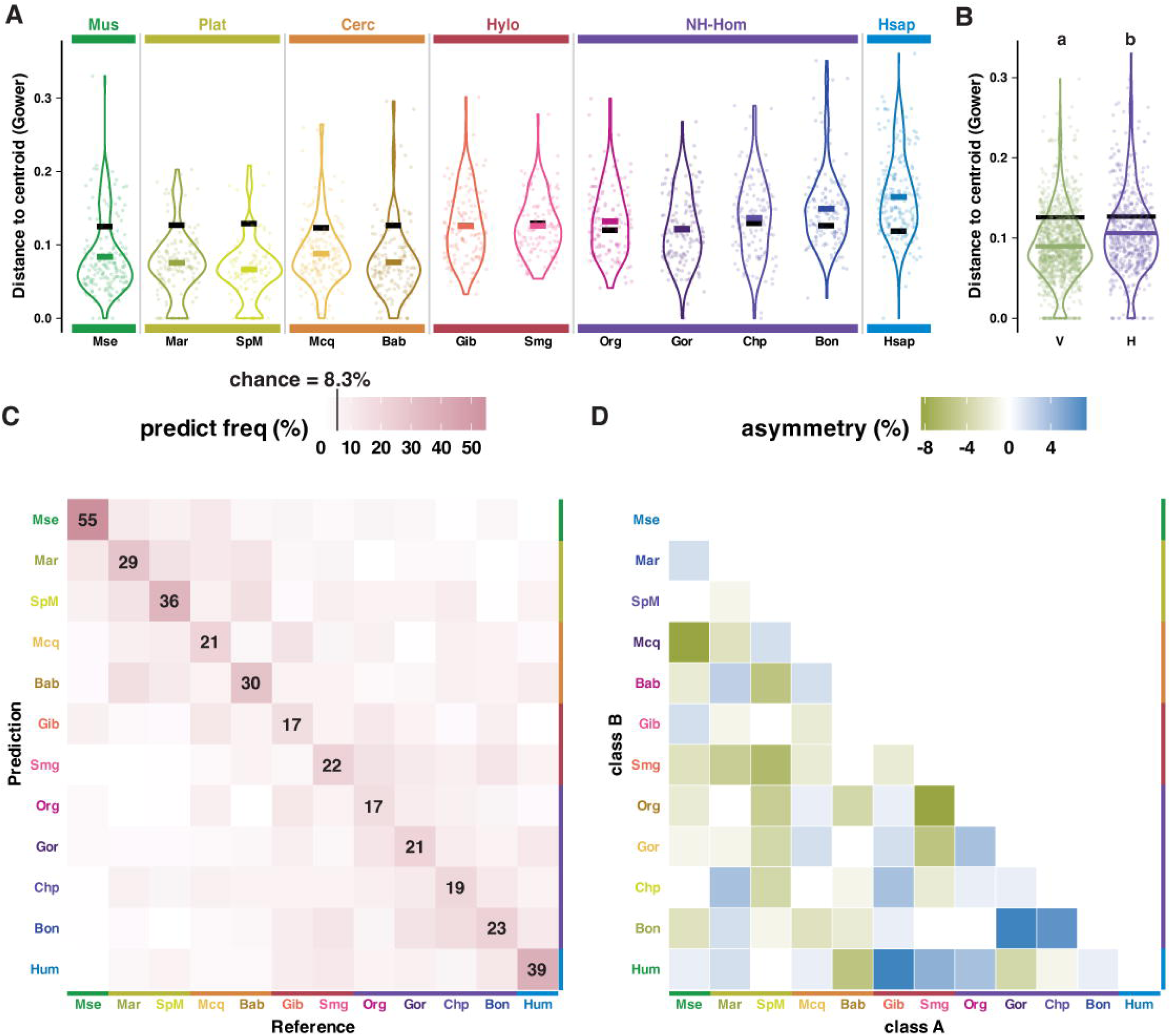
Species-level PC high dimensional dispersion and classification accuracy. A) Gower distance of each PC to the species centroid in high-dimensional space. B) Gower distance of cells to the centroid of either vertically or horizontally oriented subpopulations. C-D) Confusion matrix and off-diagonal asymmetry heatmaps for random forest classifications of PCs to each species. With 12 species tested, chance prediction is 8.3%.

**Figure S8:**
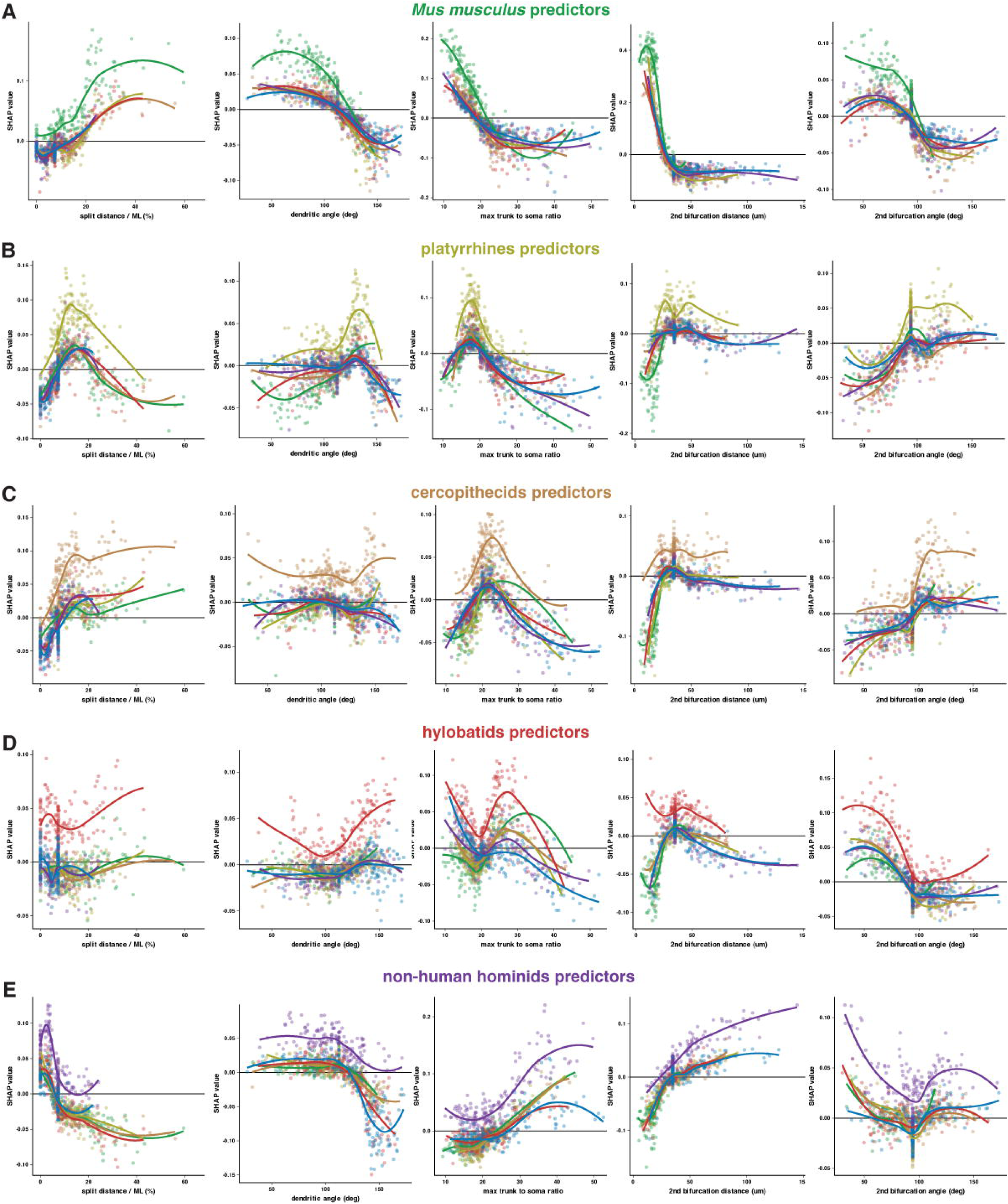
Top morphological features driving random forest classifier class-specific decisions. Shapley additive explanations analysis of input morphological variable contributions to random forest classifier decisions to predict clade identity—*Mus musculus* (A), platyrrhines (B), cercopithecids (C), hylobatids (D), and non-human hominids (E)—compared with the original values of each PC. SHAP value is positive for PCs where the variable value supported selection of the target clade and negative for values that rejected the target clade. A subset of PCs cluster at a single value because classification and the SHAP analysis were performed following median imputation to fill absent values.

**Figure S9:**
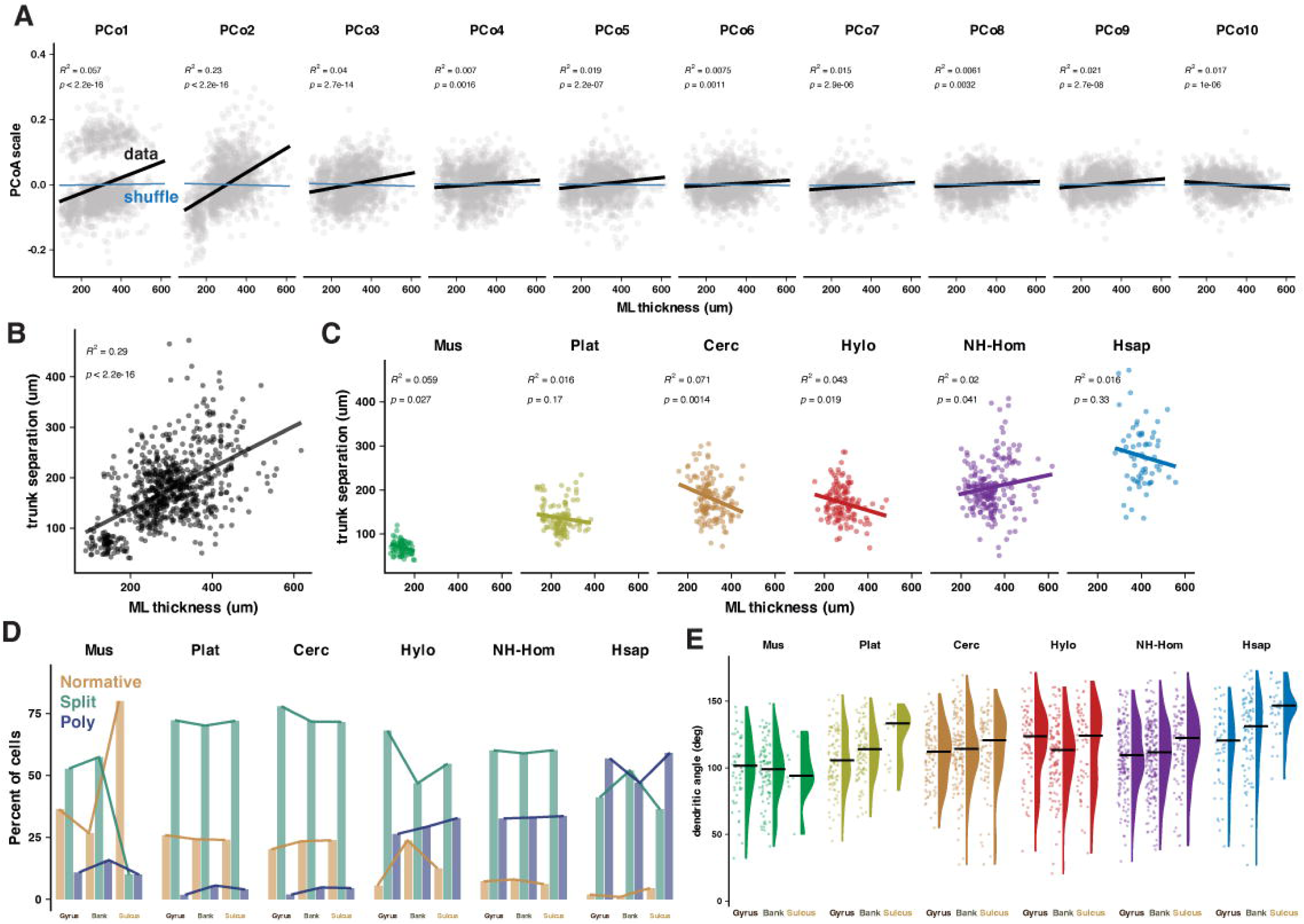
Cerebellar tissue scaling doesn’t explain PCoA axes. A) Linear regressions (black, data in grey) between ML thickness and each axis of the principal coordinate analysis of gower distances and shuffled data (blue). B-C) The relationship between trunk separation and ML thickness exemplifies that population level relationships can belie a lack of effect or even inverse relationships (Simpson’s paradox) within a clade. H) Distribution of PC morphological categories by clade and foliar region. I) Dendritic angle by foliar region.

**Figure S10:**
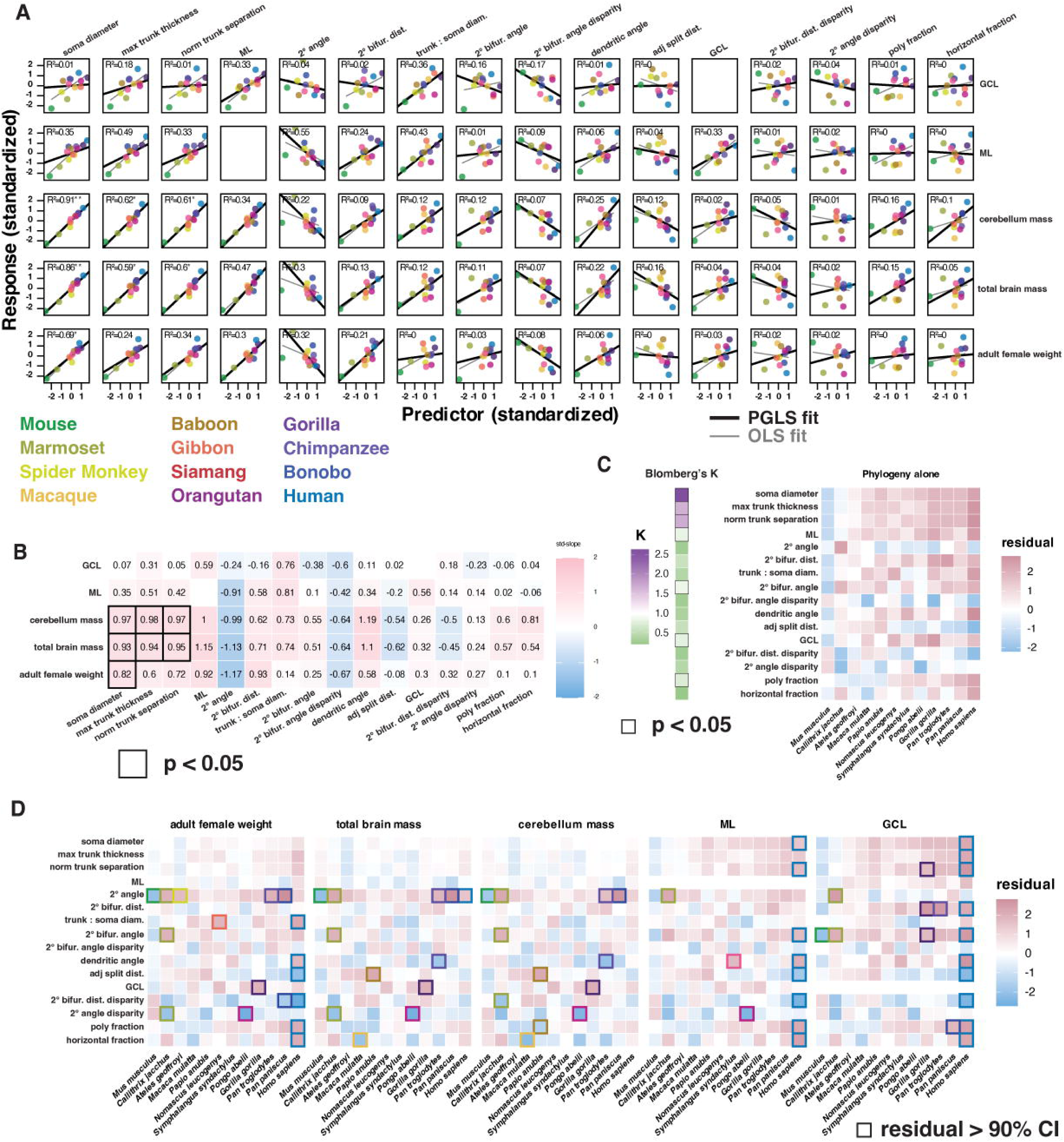
PC morphology modestly correlates with brain size but not body and local cortical scale. A) Matrix of species median morphological parameters (rows) regressed to allometric predictors (columns) using fit lines of ordinary least squares (OLS, grey) or phylogenetic generalized least squares (PGLS, black) models. Body and brain size predictors were log scaled. All predictors and parameters were standardized by centering them to the population mean and dividing by the population standard deviation. Matrix rows are sorted top to bottom by highest to lowest average R^2^ value. B) Tile matrix of standardized slope values from PGLS analysis. Sorting as in (A). Tiles with borders are statistically significant (p < 0.05 following Benjamini-Hochberg correction for multiple comparisons). C) Blomberg’s K analysis (left) of regressing each morphological variable to phylogenetic covariance under pure Brownian motion alone. Tile borders as in (B). Sorting as in (A). D) Species residuals from each PGLS regression model. Tile borders indicate residual values exceeding the 90^th^ percentile of expected residual values from pure Brownian motion generated null data. Human residuals often deviate substantially from the allometric fit line for body mass and local cortical thickness, but not brain mass. Sorting as in (A).

## Methods

### Subjects

#### Specimens

Human cerebellar tissue was collected from three unembalmed donor bodies, one provided by the Anatomical Gift Association of Illinois (AGAI) and two by the New York Brain Bank via Dr. Phyllis Faust of Columbia University. During life, all study subjects signed informed consents. Non-human primate tissue was sourced in partnership with the Association of Zoos and Aquariums (AZA). Chimpanzee cerebellar tissue was obtained from the National Chimpanzee Brain Resource (www.chimpanzeebrain.org). One macaque specimen was kindly provided by Dr. Sliman Bensmaia and one marmoset specimen was kindly provided by Dr. Nicho Hatsopoulos, both in the Department of Organismal Biology and Anatomy and the Neuroscience Institute at the University of Chicago. The remaining non-human primate tissue was provided by Dr. Chet Sherwood at George Washington University. To the extent possible, primate specimens were selected for matching adult age, covering both sexes, the absence of cerebellar complications in life or at death, and short duration in fixative for storage to avoid tissue shrinkage and improve immunolabeling quality. Table S1 provides additional demographic information about all specimens. Mouse tissue preparation was in accordance with the University of Chicago Animal Care and Use Committee guidelines. We used adult wildtype C57BL/6J mice housed on a 12hr light/dark cycle. Animals of either sex were used in all experiments and no sex differences were observed in any reported measures.

#### Allometric and phylogenetic data

Species’ mean adult brain and body size were taken as an average of data for the specimens in this study when available and otherwise compiled from numerous sources with age and sex matched data where tissue shrinkage from long term storage was accounted for as best as possible^104–110^. Euarchontoglires phylogenetic tree data was obtained from an assembled public knowledge base (Kumar et al., 2022; www.TimeTree.org).

### Immunohistochemistry

#### Human tissue preparation

Preparation of one hemisphere from the unembalmed specimens followed the same procedure as previously reported^36^. In brief, whole cerebellums were fixed in 4% paraformaldehyde (PFA) for one week after being obtained. Specimens were sectioned by hand in the parasagittal axis to obtain 3-5mm blocks that were sometimes cut transversely and further fixed for 2-4 days. Blocks were rinsed in 0.01M phosphate buffer saline (PBS) and sliced at 50µm in the parasagittal plane with a vibratome (Leica VT-1000S). Slices selected for immunolabeling were transferred to a clear tray, placed over a broad-spectrum LED array, covered with a reflective aluminum foil lid, and photobleached at 4 for 3-4 days. This reduced the strong autofluorescence in the green channel before immunolabeling.

#### Non-human primate tissue preparation

Some individuals (all macaques, all marmosets, one baboon) were anesthetized and perfused with 4% PFA. Brains were then stored in 0.01M PBS with 0.1% sodium azide. Brain tissue from the remaining specimens was removed post-mortem and immersion fixed with 4% PFA and stored in 0.01M PBS with 0.1% sodium azide. Cerebellar specimens were sectioned by hand in the parasagittal axis to obtain 3-5mm blocks that were sometimes cut transversely. Blocks were rinsed in 0.01M PBS and sliced at 50µm in the parasagittal plane with a vibratome (Leica VT-1000S).

#### Mouse tissue preparation

Mice were anesthetized with ketamine/xylazine (100 and 10 mg/kg) and perfused with cold 4% PFA. Cerebellums were removed and incubated for 2 hr in 4% PFA at 4°C and then overnight in 30% sucrose in 0.1 M PB at 4°C (until the tissue sank from the surface). The tissue was then rinsed briefly in 0.1 M PB, dried and blocked, submerged in OCT medium, flash- frozen, and then sliced at 50 μm using a cryostat microtome (CM 3050S, Leica).

#### Calbindin immunohistochemistry

Tissue was washed in 50mM Glycine in 0.01M PBS for 2hrs at 4°C and incubated in 20mM Sodium Citrate in 0.01M PBS at 80-90°C using a heated water bath for 40min. After cooling to room temperature (RT), tissue was washed in PBS with 0.5% Tween-20 (PBS-Tween) for 3x15min. Next, slices were permeabilized at RT in 0.01M PBS containing 0.025% Triton-X (PBS-TX) for 1hr and then in PBS-Tween and 200um Glycine for 15 min. Blocking was done with PBS-Tween containing 10% normal donkey serum (NDS) and 5% bovine serum albumin (BSA) for 1hr at RT followed by incubation in guinea pig anti-calbindin primary antibody (1:1000; Synaptic Systems Cat# 214 004, RRID:AB_10550535) solution overnight (18-20hrs) at 4°C with 1% normal donkey serum in PBS-Tween. After 3x10min washes in PBS-Tween at RT, slices were incubated in donkey anti-guinea pig Cy3 secondary antibody (1:200; Jackson ImmunoResearch Labs Cat# 706-165-148, RRID:AB_2340460) for 2hrs at 4°C with 1% NDS in PBS-Tween. Finally, slices were washed in PBS-Tween for 3x10min, mounted and coverslipped with Vectashield (Vector Laboratories, Inc.), and allowed to set overnight before visualization. For mouse tissue, glycine incubation lasted only 1 hr and the heated sodium citrate incubation lasted only 20 min.

### Confocal Imaging

#### Confocal Imaging for Population measurements

Following immunolabeling, we identified 5-10 representative regions across gyrus bank and sulcus sub-regions of Crus 1 that contained as many PCs as possible. We then collected z-stack images throughout the whole section thickness at 10x (Zeiss Achroplan 0.25NA, air). Images were then marked for cell type categories using the ObjectJ package in ImageJ and measured for the below described morphological parameters using standard measurement tools in ImageJ.

### Cell Classification and Measurement Definitions

#### Purkinje cell morphological category definitions and criteria

Criteria for morphology category definitions were the same as previously^11,36^ but we reiterate the full description here for clarity. PCs were deemed ***Normative*** if they had the following features: 1) a single trunk emerging from the soma, and 2) either no bifurcation of the primary trunk within two soma distances (2x the diameter of the soma) or a highly asymmetrical bifurcation where the smaller branch did not project in the parasagittal axis more than ∼5x the somatic diameter from the main dendritic compartment. PCs were defined as ***Split*** if they had the following features: 1) a single trunk emerging from the soma, and 2) either symmetrical bifurcation of the primary trunk within two soma distances or an asymmetrical bifurcation within two soma distances where the smaller branch projected more than ∼5x the somatic diameter from the main dendritic compartment and thus reached prominence by its overall length and sub-branching. PCs were defined as ***Poly*** if they had more than one trunk emerging from the soma regardless of relative size. All PCs were further subdivided into Vertical or Horizontal ramification patterns. PCs were defined as ***Horizontal*** if one primary dendrite ramified parallel (within 30 degrees) with the PC layer for

∼7x the somatic diameter or both primary dendrites ramified in opposing directions parallel with the PC layer for 4x the somatic diameter. Otherwise, the cell was defined as ***Vertical***.

#### Foliar sub-region category definitions and measurement criteria

Criteria for foliar category definitions were the same as previously^11,36^ but we reiterate the full description here for clarity. Purkinje cell locations were defined as either Gyrus, intermediate Gyrus, Bank, intermediate Sulcus or Sulcus based on the relative expansion/compression of the granule cell (GCL)/molecular layers (ML) in the parasagittal axis. ***Gyrus*** was defined as a region where the total parasagittal length of the pial surface exceeded that of the border between the GCL and the white matter. Intermediate Gyrus was defined as a region where the pial surface expanded relative to the GCL, but there was no obvious extension of white matter into the structure. ***Bank*** was defined as regions where those two lengths were approximately equal, such that neither layer of the cortex was compressed or expanded relative to the other. Intermediate sulcus was defined as regions where the ML contracted, but without gyrus structures around it. Finally, ***Sulcus*** was defined as regions where the total parasagittal length of the pial surface was less than that of the border between the GCL and the white matter.

GCL thickness was measured using the shortest line from where the PC soma abuts the GCL to where the GCL abuts the white matter. ML thickness was measured using the shortest line from where the PC soma abuts the ML to the pial surface.

#### Parametric Measurement Definitions

Measurement parameters are defined and illustrated in Figure S2. Measurements were taken only if the relevant parameter was visibly intact for a given PC. Species identity was redacted from all images during measurements and the measuring experimenter was blind to species identity. All measurements were performed in ImageJ.

##### Soma circumference, diameter, and area

A polygon was drawn around the edge of the PC soma and measured for area and circumference. Diameter was approximated with circumference/.

##### Split distance

Split distance was measured only for cells with a single primary dendrite using a segmented line along the primary dendrite, beginning at the somatic apex where the primary dendrite extrudes from the soma and ending at the primary dendrite’s first approximately symmetrical bifurcation. A segmented line was used for measuring distances to follow curvature in the dendrite. As such, distances represent dendritic length, not the Euclidean distance between origin and tip.

##### Secondary bifurcation distance

*For Normative and Split cells:* The segmented-line length from the first dendritic bifurcation to the next symmetrical bifurcation of each daughter branch. *For Poly cells:* The segmented-line length from the somatic extrusion of each primary dendrite to its first symmetrical bifurcation. *Dendrite disparity:* The minimum of a given PC’s secondary bifurcation values divided by the maximum.

##### Trunk thickness

*For Normative and Split cells:* The straight-line diameter of a PC’s primary dendrite, measured where the base of the trunk emerges from the continuing curvature of the soma. *For Poly cells:* The widths of each primary dendrite, measured where they emerge from the soma. *Dendrite disparity (Poly only):* The minimum of a Poly PC’s trunk thickness values divided by the maximum.

##### Dendrite separation

The straight-line distance between dendritic compartment centroids measured parallel with the PCL/Pial surface and at half the thickness of the ML. The centroid was visually estimated to be the center of mass of the two sides of the arbor. This often, but not always tracked with the bisector of the secondary bifurcation angle, which tended to define the subsequent architecture of the distal tree.

##### Dendritic angle

*For Normative and Split cells:* The branching angle at the first bifurcation of the primary dendrite. *For Poly cells:* The angle between the predominant directions formed by each trunk up to the first symmetric bifurcation.

##### Secondary bifurcation angle

The branching angle of each secondary dendritic bifurcation. *Dendrite disparity:* The minimum of a given PC’s secondary bifurcation angles divided by the maximum.

##### Secondary angle

The angle created by the intersection of the trajectory from the primary bifurcation point to the secondary bifurcation point and the bisector of the secondary bifurcation angle. *Dendrite disparity:* The minimum of a given PC’s secondary angles divided by the maximum.

##### Arc complete

*For Normative and Split cells:* The arc distance from one side of the primary dendrite to the other. For cells with one trunk, this was effectively the circumference minus the trunk thickness. *For Poly cells:* The arc distance from the GCL side of one trunk to the other along the underside of the soma.

##### Arc to axon

The arc distance along the somatic circumference from the axon to the nearest primary dendrite.

### Parameter Analysis

#### Morphospace analysis

Morphospace was defined by the set of input variables, which included: Number of trunks, number of compartments, Split Distance normalized to ML thickness, Primary dendritic angle, Secondary bifurcation distance, mean secondary angle, mean secondary bifurcation angle, maximum trunk thickness / soma diameter, compartment centroid distance normalized to ML thickness, Disparity between secondary angles, disparity between secondary bifurcation distances, disparity between secondary bifurcation angles, and disparity between multi-dendritic PC trunk thicknesses. Distance matrices were calculated with Gower distances, instead of Euclidean, as our morphological parameters have mixed statistical structure (continuous vs discrete, bounded vs unbounded) and often lack values owing to tissue quality (biological missingness) and variables not always applying (structural missingness). Using the distance matrix, we performed principal coordinate analysis (PCoA) dimensionality reduction. We preserved row identity, allowing us to match identifying information for each cell to the extracted PCoA axes. This also allowed us to assess multivariate dispersion of cells across different identity groups (clade or morphological category). Shuffled data consisted of an all-to-all shuffle of each input variable while cell identities (species, clade, location, etc) were held constant.

#### Feature variance calculations

Variability of each morphological parameter was quantified using complementary dispersion metrics. For continuous variables, we calculated a robust coefficient of variation defined as the median absolute deviation divided by the median, which reduces sensitivity to outliers. Discrete traits were analyzed separately using Shannon entropy to measure categorical diversity. To assess morphospace occupation, we computed the fraction of the global range for each variable spanned by each group. To enable comparison across variables with different units, values were additionally rescaled to the interval [0,1] and dispersion quantified as the standard deviation of normalized values. Missing observations were excluded from calculations. Metrics were computed globally and within clades.

##### Robust coefficient of variation

Shannon entropy

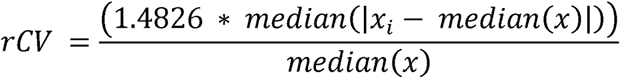

##### Shannon entropy

Let *p_k_*be the proportion of observations in category *k*.

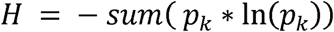

where

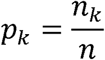

##### Fraction of range occupied

Let

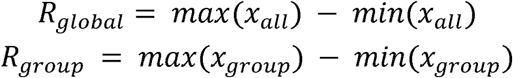

Then

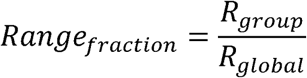

##### Normalized standard deviation

Normalized observation *x_i_* is *x_i,norm_*

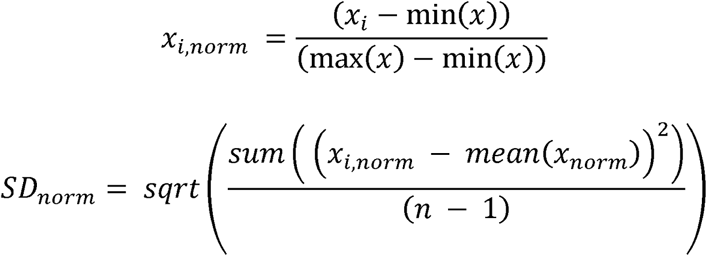

#### Classifier and post-hoc SHAP analysis of feature contributions

To classify PCs by clade, we trained a random forest classifier using the same morphospace variables above. To account for structural missingness (e.g. Poly PCs do not have a split distance), we added binary indicator variables to indicate if each parameter had a measurement. To obtain a complete input matrix despite many missing values, we filled gaps without biasing toward any one clade profile through median imputation. To prevent the model from favoring morphological statistics from one clade, we randomly down sampled without replacement PCs from each clade to the number of values present in the lowest sampled clade (166 mouse cells) or species (108 Spider Monkey cells). Any predictors with near-zero variance were removed prior to model fitting. The classifier model was built using 5-fold cross-validation with automatic parameter tuning optimization. Prediction accuracy was summarized using recall-normalized confusion matrices, which express prediction frequency within each true class.

Feature contributions to the classifier model predictions of each clade were quantified post-hoc by using a descriptive SHAP (Shapley Additive Explanations) analysis. SHAP was computed with a Monte-Carlo sampling approach (50 permutations per observation) to estimate each variable’s marginal contribution to the predicted probability of a single target clade at a time. We applied a finite-sample bias correction as the dataset is relatively small, though some features of a finite bias, such as favoring high variance variables or dependent features, are less likely to play a role as separate analyses of parameter variance (above) already identified almost all input variables as having high entropy and little covariation. Global feature importance was quantified as the mean absolute SHAP value, while directionality (the observation’s tendency to push toward or away from the target class prediction) was assessed using mean signed SHAP values. SHAP values are given as contribution to probability as opposed to log-odds or internal scoring.

#### Statistics

Statistical analysis was carried out using R (v4.4.2). A Wilcoxon rank sum test was used for two group comparisons. Group comparisons were conducted using pairwise Kruskal-Wallace tests with FDR correction for multiple comparisons followed by a Dunn post-hoc test to detect separable groups. We used a Pearson’s Chi-squared test for independence to assess contingency tables from data with two non-continuous variables. PCoA axis eigenvalues were compared to a shuffled distribution via sequential Student’s t-tests applied until failure to reject the null.

## Notes

### Competing Interest Statement

The authors have declared no competing interest.

### Funding

- National Institutes of Health (NINDS): R21NS124217
- National Institutes of Health (NINDS): F31NS129256

